# Retracing the evolutionary trajectory of adenine base editors using theoretical approaches

**DOI:** 10.1101/2020.12.24.424366

**Authors:** Kartik L. Rallapalli, Brodie L. Ranzau, Kaushik R. Ganapathy, Alexis C. Komor, Francesco Paesani

**Author notes:** Contributed equally to this work (Brodie L. Ranzau, Kaushik R. Ganapathy).

## Abstract

Adenine base editors (ABEs) have been subjected to multiple rounds of mutagenesis with the goal of optimizing their function as efficient and precise genome editing agents. Despite this ever-increasing data accumulation of the effects that these mutations have on the activity of ABEs, the molecular mechanisms defining these changes in activity remain to be elucidated. In this study, we provide a systematic interpretation of the nature of these mutations using an entropy-based classification model that relies on evolutionary data from extant protein sequences. Using this model in conjunction with experimental analyses, we identify two previously reported mutations that form an epistatic pair in the RNA-editing functional landscape of ABEs. Molecular dynamics simulations reveal the atomistic details of how these two mutations affect substrate-binding and catalytic activity, via both individual and cooperative effects, hence providing insights into the mechanisms through which these two mutations are epistatically coupled.

## Introduction

The ability to introduce A•T to G•C base pair conversion in the genetic code of an organism, in a precise, efficient, and programmable manner, has the potential to correct almost 60% of known pathogenic point mutations in human beings. ^1^ Targeted A•T to G•C conversions have recently been realized through the development of adenine base editors (ABEs).^2^ ABEs consist of two subunits: a catalytically-impaired Cas9 (Cas9n), which serves as a programmable DNA-targeting module, and an engineered variant of a tRNA adenosine deaminase enzyme (TadA*),^3^ which serves as the single-stranded DNA (ssDNA)-editing module and enables the hydrolytic deamination of targeted adenosines (A) into inosines (I). Inosine is subsequently converted into guanosine (G) by the DNA repair and replication machinery, completing the A•T to G•C base pair conversion by ABEs (Figure 1A).

**Figure 1:**
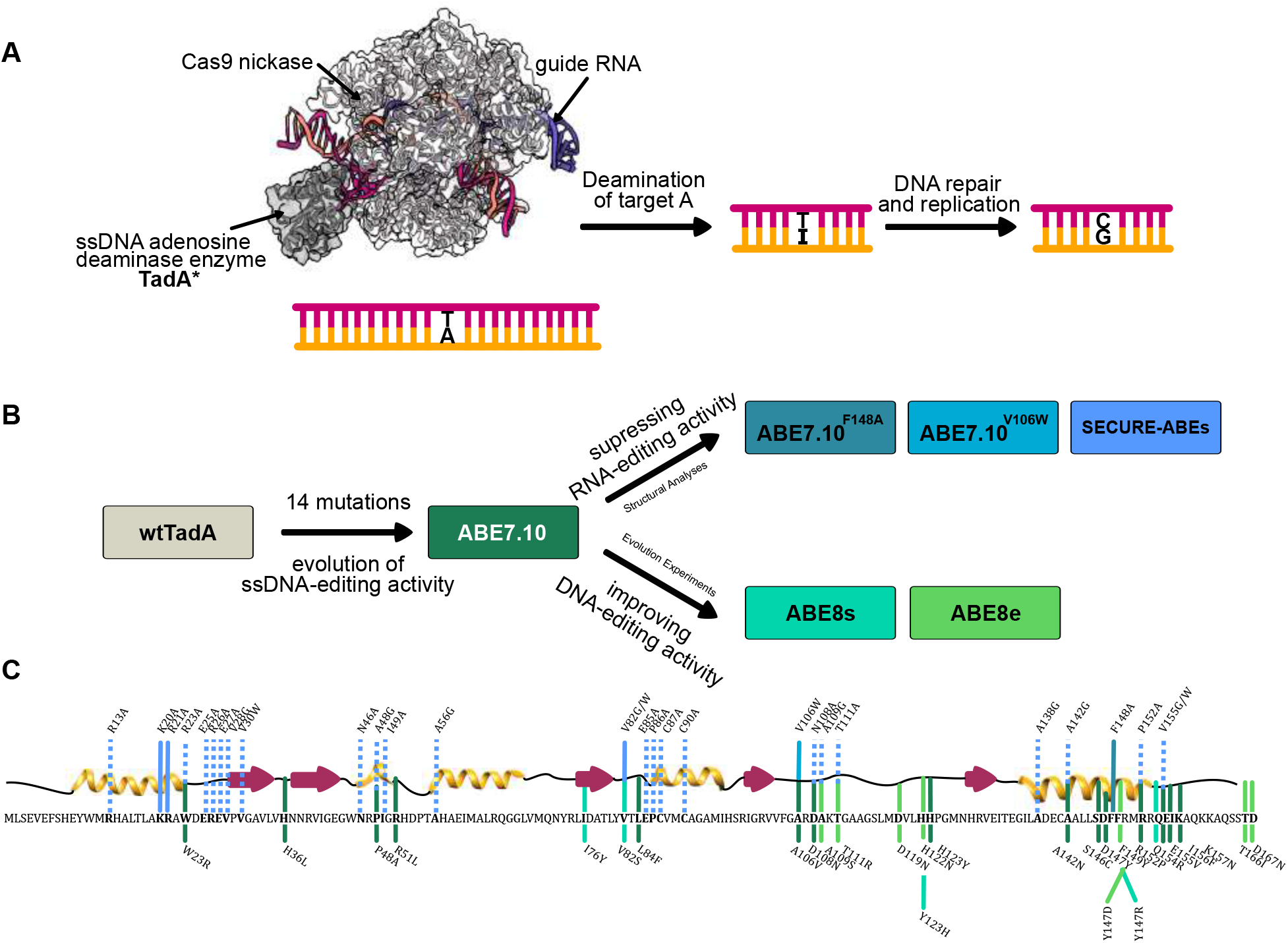
(A) Schematic representation of base editing by ABEs (PDB ID: 6VPC).^18^ The binding of Cas9n to the target genomic locus unwinds the DNA double helix and exposes a small region of ssDNA. TadA* hydrolytically deaminates the adenine (**A**) to form inosine (**I**) which is subsequently converted to guanine (**G**) by cellular DNA repair and replication machinery. (B) Engineering efforts in the field to generate and improve upon ABEs, starting from *E. coli* wtTadA. (C) Primary and secondary structure of *E. coli* wtTadA with key mutations indicated. The line colors correspond to colors shown in (B), indicating the ABE version in which these mutations were identified. Solid lines are mutations that were incorporated into final ABEs constructs, while dashed lines are mutations that were experimentally tested in previous work, but not incorporated into final ABE constructs.

ABEs continue to remain a focal point of interest for the genome editing community, not only because of their potential as therapeutic agents^4–9^ but also because of the remarkable scientific effort that went into their development. Extensive protein engineering and evolutionary methods were employed to impart ssDNA-editing capabilities onto an RNA-editing enzyme, wild-type *E. coli* TadA (wtTadA), resulting in the seminal ABE7.10 base editor.^2^

Although the mutations that gave rise to the original ABE7.10 construct successfully imparted ssDNA-editing capability onto TadA*, they did not suppress the inherent RNA-editing activity of TadA*. It was subsequently demonstrated that ABE7.10 induces considerable gRNA-independent off-target RNA-editing throughout the transcriptome.^10–12^

Since the development of ABE7.10, major efforts have been devoted to its further evolvement on two separate fronts (Figure 1B). First, additional rounds of directed evolution were employed to increase the on-target ssDNA-editing activity by TadA, resulting in ABE8.20^13^ and ABE8e.^14^ Second, structural analyses of the TadA–RNA complex followed by rational engineering was employed to decrease the off-target RNA-editing activity by TadA, resulting in ABE7.10^*F* 148*A*^,^15^ ABE7.10^*V* 106*W*^,^16^ and SECURE-ABEs.^17^

Due to the lack of naturally occurring ssDNA-editing enzymes, the expansion of the existing base editing repertoire would inevitably require evolution and engineering strategies on new enzymes to first introduce ssDNA-editing activity, followed by structure-based redesign to abrogate inherent RNA-editing activity, analogous to those used in the development of ABEs. The success of structure-based protein engineering efforts are highly dependent on the availability of appropriate X-ray or cryo-EM structures of the protein–RNA complex. Even when structural data are available to guide this process, most mutations, especially those concentrated near the active site of the enzyme, are likely to have detrimental effects on the enzyme’s function.^16,17,19^ Hence, it is important to fully understand the features that are essential for the native RNA-editing function of TadA*, and how certain mutations can affect changes to its substrate binding and catalytic mechanism.

To date, many studies have mutated ABEs to manipulate its DNA- and RNA-editing abilities, producing a large amount of experimental data associated with mutations at 45 of the 167 residue sites of TadA* (Figure 1B and C). To gain fundamental insights into ABE’s editing activity from this ever-expanding pool of mutational information and guide future efforts in the development of new base editors, we have carried out a systematic data-driven computational study combined with experimental assays to better understand, in atomistic detail, the effects of individual mutations on the activity of TadA*.

## Results and Discussion

### Dataset Curation and Sequence Entropy Calculation

The principal tenet of biochemistry is that the primary sequence of amino acids comprising a protein dictates its three-dimensional molecular structure, which then determines its biological function.^20^ To date, most ABE engineering efforts have relied on the second and third tiers of this tenet, in the form of structural analyses^15–17^ followed by site-directed mutagenesis and experimental measurements of the resulting functional properties of TadA (second tier) or directed evolution where TadA is randomly mutated and functional variants are identified through a selection scheme (third tier)^2,13,14^ (Figure 1B). Due to the time- and resource-intensive nature of these second and third tier methods, we decided to begin our investigations by focusing instead on the first tier of this tenet, that is, on investigating how the primary sequence of TadA can be used to rationalize the effects that individual mutations, identified experimentally, have on the native function of TadA* (i.e. RNA-editing activity) (Figure 1C).

With the expansion of reliable protein sequence databases, ^21,22^ the statistical analyses of protein homologs have already enabled the successful prediction of mutational effects on the function of several enzymes,^23^ including cytosine base editors.^24^ For our sequence-based analyses of the ABE mutations, we used the amino acid sequence of *E. coli* wtTadA^3^ as our query for a BLAST search^25^ of its extant homologs in the SWISSPROT database, ^22^ which generated a dataset of 75 homologs. However, as our primary focus was to identify residues essential for the function of TadA on its native RNA substrate, we filtered out distant homologs using stringent percentage identity and coverage length cutoff values (Figure 2A). This filtering resulted in a more focused dataset as it removed functionally distinct and redundant sequences from our initial BLAST search. Despite reducing the size of the dataset considerably (to 35 homologs), this filtered dataset still captures the diversity of our initial unfiltered dataset (Figure 2A).

**Figure 2:**
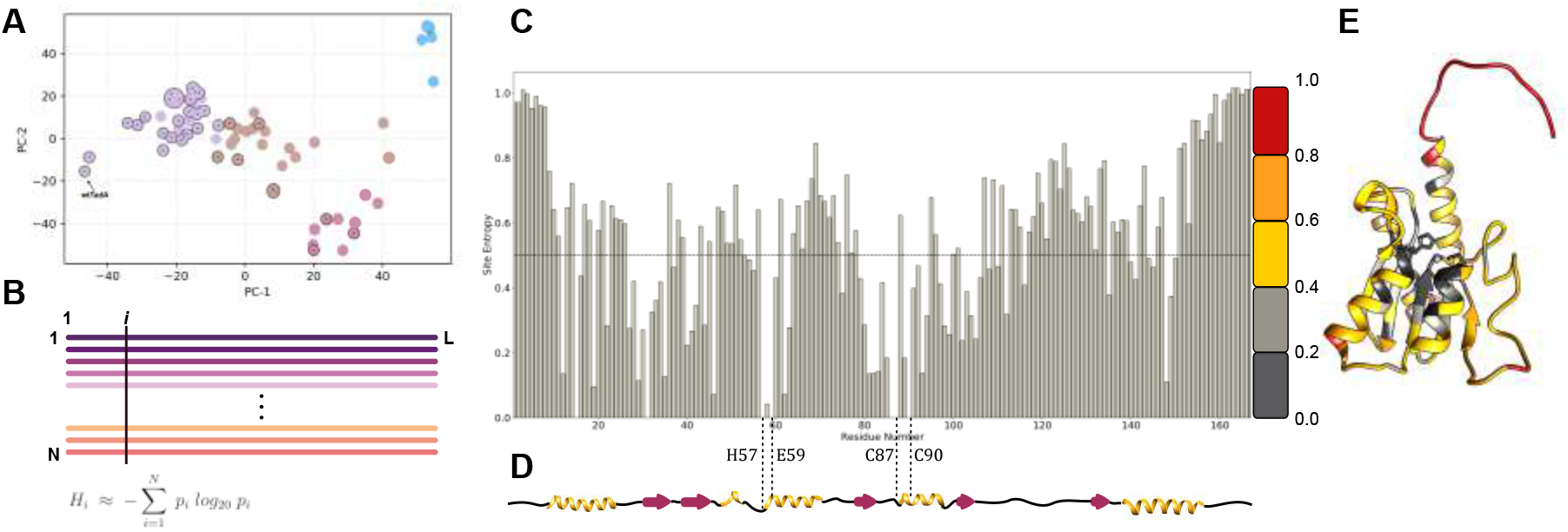
(A) The top two principal components of the pairwise sequence distance matrix of extant homologs comprising the filtered (indicated with circles and dots) and unfiltered datasets. Based on the similarity of the sequences, the dataset is clustered into 4 separate sets, colored purple, brown, red, and blue. The sequences in the filtered dataset are highlighted in each cluster. (B) Multiple sequence alignment of extant homologs of wtTadA to calculate the statistical probability of occurrence of individual amino acids at residue site *i* (*p*_*i*_). This is subsequently used to assign a conservation score to site *i*, using Shannon’s definition of information entropy (*H*_*i*_) Equation 1. (C) Information entropy of individual residue sites of the wtTadA query, with its secondary structure elements mapped below in (D). (E) Entropy values mapped on to the three-dimensional structure of *E. coli* TadA using a color gradient to signify conserved residues and mutational hotspots.

To visualize the sequence space captured by our unfiltered and filtered datasets, and highlight relationships among these wtTadA homologs, we performed a dimensionality reduction of the dataset using principal component analysis (PCA). This allows us to project the hyper-dimensional sequence space associated with the homologs onto two dimensions, while still preserving the relationships among the various homologs. By partitioning the dataset into four representative clusters (Figure S1), the outcome of filtering becomes more apparent as each cluster consists of functionally similar homologs. These clusters are represented by different colors (purple, brown, red, and blue) in Figure 2A. From this analysis, it can be observed that this filtered dataset indeed captures the overall diversity of the unfiltered dataset, as three of the four clusters are represented. The purple cluster (containing the query sequence) consists primarily of TadAs and their eukaryotic equivalents, ADAT2s, and hence has the greatest number of filtered sequences, 26 of the 35 filtered sequences. The next most populated cluster, the brown cluster, consists of 22 sequences in the unfiltered set, which results in 6 sequences after filtering. Only three out of eleven sequences were selected from the red cluster, with two of these sequences belonging to the cytidine deaminase superfamily (and are on average 50% similar to the query sequence) and the third sequence corresponding to a guanine deaminase. Given the distance between the blue cluster and the query sequence (i.e. the lack of similarity between these sequences, which mostly represents the catalytically inactive Tad3 and ADAT3 proteins), it is not surprising that no sequences were selected from this cluster upon the implementation of our filters. It is important to note that our filtered dataset consists entirely of RNA-editing enzymes, demonstrating the effectiveness of our filters. We, therefore, reasoned that any primary sequence analyses of our filtered dataset would be highly biased towards illuminating aspects of RNA-editing activity by the wtTadA (Figure S1B).

Having obtained a reliable dataset of extant TadA homologs, we next sought to quantify the evolutionary conservation and functional importance of individual residues of wtTadA. An extensively studied and widely used approach to address this problem is the evaluation of information theory-inspired sequence entropy scores (Figure 2B). ^26–29^ Within this approach, the sequence entropy for each residue site, *i*, is defined as:

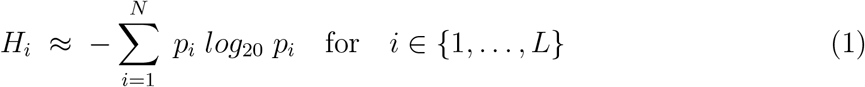

where *p*_*i*_ refers to the statistical probability of having a particular amino acid at site *i*. Thus, the value of *H*_*i*_ ranges between 0 and 1, with an entropy value of 0 indicating that the site has only one unique amino acid represented within the dataset (suggesting that the site is highly conserved from an evolutionary standpoint), and an entropy value of 1 indicating that the site has every possible amino acid represented within the dataset (suggesting that such a site is naturally more tolerant to mutations).

Applying Equation 1 to the filtered dataset of TadA homologs (Figure 2A), we calculated the site entropy for the entire wtTadA sequence (Figure 2C). The active site of wtTadA consists of a zinc ion tetrahedrally coordinated by Cys^87^, Cys^90^, His^57^, and a water molecule. This water molecule is activated for deamination reaction by the highly conserved Glu^59^ residue. Consistent with the importance of these residues for the canonical RNA-editing activity of TadA, we observed *H*_*i*_ = 0 for these four active site residues. This active site is further stabilized by a *β*-sheet core, and the entropy values for 24 of the 38 core residues are also low (*H*_*i*_ ∈ {0.0, 0.4}). The surface-exposed residues have relatively higher values of *H*_*i*_, with the C- and N- terminal residues having the highest values (*H*_*i*_ > 0.4) (Figure 2D). By mapping these entropy scores onto the structure of wtTadA^30^ (Figure 2E), the correlation between the entropy values and the three-dimensional structure of TadA is clearly apparent. Thus this sequence-based entropy model is capable of representing the structural information encoded by the amino acid sequence of wtTadA.

### Sequence Entropy as a Binary Classifier of TadA* Function

Building upon these results, we used the sequence-based entropy model to rationalize the role played by different amino-acid mutations that have been experimentally shown to modulate the function (i.e. RNA-editing activity) of ABEs (Figure 1B and C). Based on the biochemical interpretation of the two extreme entropy values, we chose *H*_*i*_ = 0.5 as an initial cutoff value to distinguish the functional implications of the entropy data obtained for the wtTadA sequence (Figure 2C) in the context of all mutations reported for the ABEs (Figure 1B and C). Within this model, we hypothesize that residue sites having *H*_*i*_ > 0.5 will either induce no change in the activity of wtTadA or, if mutated appropriately, can have a favorable impact on the native activity (i.e., RNA-editing activity) of wtTadA. Conversely, sites with *H*_*i*_ ≤ 0.5 are predicted to have adverse effects on the canonical RNA-editing activity of wtTadA.^15–17^ It should be noted that, since our dataset comprises only RNA-editing enzymes, we are primarily referring to the impact that individual mutations have on the native function of the wtTadA sequence (SI sequences and Figure S1B). However, given the vast amount of experimental data available regarding the mutations that impact the ssDNA-editing efficiency of TadA*, we were interested in assessing the performance of the entropy-based model on these mutations as well. Hence, mutations that either increase the ssDNA-editing ability or had no negative impact on the RNA-editing activity of ABEs (as discovered using either directed evolution^2,13,14^ or site-directed mutagenesis^17^) are deemed to be correctly classified using our information entropy-based model if their *H*_*i*_ value is greater than 0.5 (Figure 3A).

**Figure 3:**
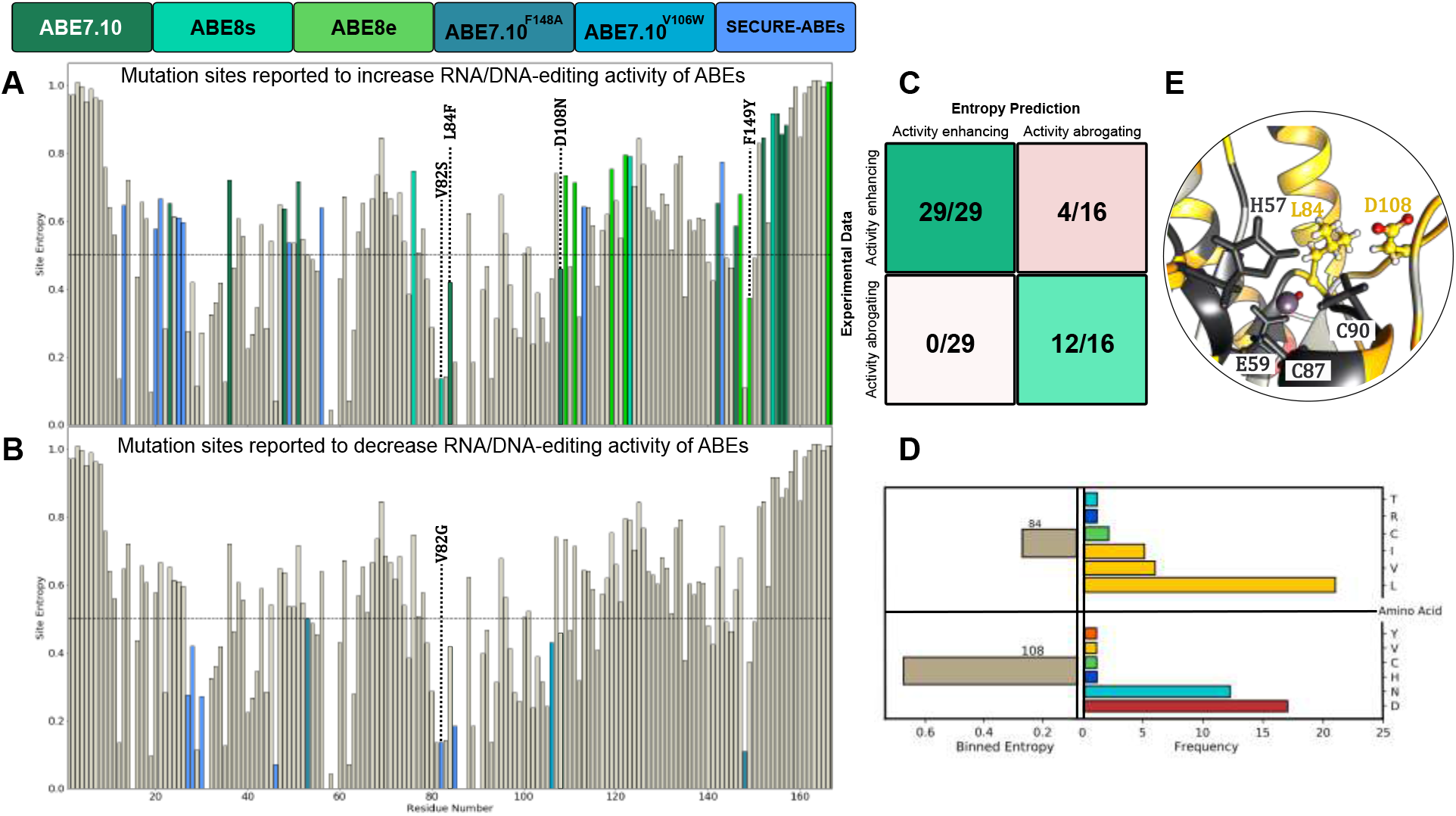
(A) Mutations reported to have beneficial or neutral effects on the RNA or DNA editing activity of the ABEs. (B) Mutations reported to have detrimental effects on the RNA or DNA editing activity of the ABEs. (C) Confusion matrix of the experimental data and the entropy-based classifier. (D) Binned entropy values and distribution of amino acids at sites 84 and 108. (E) Local environment of 84 and 108 residues in the wtTadA structure.

To quantify the performance of the sequence-based entropy model, we computed the confusion matrix where each prediction (Figure 3A and B) is validated against the corresponding experimental editing outcome for 45 total mutations^2,13–17^ (Figure 3C). By construction, the diagonal elements of the confusion matrix thus correspond to correct predictions, while the off-diagonal elements indicate incorrect predictions. Despite being entirely derived from the information content of amino-acid sequences contained in a highly biased RNA-editing dataset, the sequence-based entropy model applied to all the reported ABE mutations exhibits a remarkable accuracy of 91.1% and an *F*_1_ score of 0.91 (Figure 3C). Specifically, the model correctly predicted all the mutations that are reported to adversely impact the native RNA-editing activity of TadA*. However, it incorrectly predicted the effects of 4 mutations, all of which correspond to residues with low entropy values that were experimentally found to increase ssDNA-editing activity. Hence, in an attempt to understand the significance of these misclassified residues, and to better understand the deficiencies of our model so as to refine our classification scheme, we sought to further analyze the amino acid distribution at these residue sites. (Figure 3A). We found that site 82 (Val in the wild-type enzyme) had ambiguous experimental data as its mutation to Gly abrogates RNA-editing in SECURE-ABEs,^17^ while its mutation to Ser results in enhanced ssDNA-editing in ABE8s. ^13^ This suggests higher predictability of the entropy classifier regarding the native RNA-editing activity of TadA* than its ssDNA-editing activity, as expected. Additionally, both sites 84 (Leu in wtTadA) and 108 (Asp in wtTadA) are associated with low entropy values, but were mutated to enhance ssDNA-editing activity during the development of the foundational ABE7.10.^2^ Similarly, a low entropy value is found for site 149 (phenylalanine in wtTadA), which was mutated to enhance DNA editing activity in ABE8e.^14^ These incorrect predictions highlight the limitations of the model and indicate that specific mutations at select low entropy sites may allow TadA to gain activity towards a different substrate (ssDNA-editing versus RNA-editing).

The D108N mutation was the critical first mutation that led to the onset of ssDNA-editing activity of TadA*.^2^ Moreover, this residue is part of a surface-exposed loop in the structure of TadA*. Hence, we would expect this residue to display high entropy. To further dissect the anomalous misclassification (*H*_*i*_ < 0.5) of site 108 through our entropy-based model, we analyzed the distribution of various amino acids at this site within our dataset (Figure 3D). To our surprise, although the mutational entropy of this site is marginally low, approximately 36% of the dataset sequences record an Asn at this site, making it the second most probable amino acid at site 108. This observation was particularly striking given the importance of the D108N mutation; it was the first mutation observed in the directed evolution of the foundational ABE7.10,^2^ and we recently discovered that reversion of this mutation in the ABE7.10 construct resulted in complete loss of ssDNA-editing activity by TadA*.^31^ It is therefore quite significant that a mutation that is so critical for imparting novel ssDNA-editing functionality onto an RNA-editing enzyme has such a high incidence in naturally-occurring TadA homologs (Figure 3D). Additionally, this indicates that in the case of TadA* evolution, the enzyme achieves activity towards DNA substrates while still retaining activity toward its native RNA targets.

Upon conducting a similar distribution analysis for site 84, which is also misclassified by the entropy-based model as a low entropy site that favorably affects ssDNA-editing, we found that while this core residue has a low sequence entropy of *H*_*i*_ = 0.416 as defined by Shannon’s entropy, 88.6% of sequences had an aliphatic amino acid (Leu, Val, or Ile) at this position, in direct contrast to its mutation to Phe as in ABE7.10 (Figure 3D). Thus, unlike the D108N mutation, the L84F mutation is a novel mutation that had not been explored by natural protein evolution.

This analysis of the distribution of the possible amino acids based on their chemical nature helps identify the types of mutations that are tolerated at various sites of the TadA* sequence (Figure S2A). Hence, we re-calculated the entropy values for wtTadA by binning amino acid residues according to their side chain classifications: polar uncharged, positively charged, negatively charged, hydrophobic-aliphatic, hydrophobic-aromatic, and special (Gly, Pro). The resulting binned entropy values (Figure S2B, C, and D) were greater than 0.5 for site 108 while still remaining lower than 0.5 for site 84 (Figure 3D). These results thus indicate that the entropy-based analysis allows not only for the quantification of the mutational propensity of individual wtTadA sites but also the characterization of the chemical properties that make mutations to a specific class of amino acids relatively more favorable. Moreover, we also speculate that residue sites having marginally low *H*_*i*_ values can in fact be mutated based on the amino acid distribution observed in its extant homologs to confer novel functionality to the enzyme (as seen for D108N mutation) or to disrupt native functionality (as seen for L84F).

### Experimental Analyses

We next sought to experimentally test our hypothesis that the conservation scores and amino acid distributions derived from the entropy-based model could be used to predict the effects of mutations on the RNA-editing activity of TadA*. It is well known that later-generation ABEs induce transcriptome-wide RNA editing, but it is unknown if this is a “carryover” activity from wtTadA being able to edit RNA sequences other than its native tRNA substrate, or if the various mutations identified through directed evolution not only enhanced the ssDNA-editing activity of TadA*, but also its nonspecific RNA-editing activity. We first tested the ability of ABE0.1 (both as monomeric and dimeric wtTadA fused to Cas9n) to introduce A-to-I edits in mRNA in a gRNA-independent manner. We transfected HEK293T cells with constructs encoding monomeric ABE0.1, dimeric ABE0.1, or heterodimeric ABE7.10 (wtTadA-TadA*-Cas9n), extracted mRNA after 36 hours, and used high throughput sequencing to quantify A-to-I editing at six different sites throughout the transcriptome that had previously been shown to be edited by ABE7.10 in a gRNA-independent manner.^17^ To our surprise, we observed >50% A-to-I RNA-editing efficiencies at all six sites by both wild-type constructs; in fact, RNA-editing activity by dimeric ABE0.1 was on average 20% higher than ABE7.10. Our entropy-based analysis suggests that non-aliphatic mutations at site 84 would diminish the RNA-editing activity of wtTadA, while certain mutations at site 108 would retain (or even enhance) the RNA-editing activity of wtTadA (Figure 3 and Figure S2). To test this hypothesis, we generated six different ABE variants - ABE1.1 (i.e. ABE0.1(D108N)), ABE0.1(L84F), and ABE1.1(L84F), and their corresponding heterodimeric constructs (i.e. wtTadA-TadA*-Cas9n), and compared their RNA-editing activities with ABE0.1 and ABE7.10 at the same six sites. Each variant was tested as a monomer and a heterodimer to account for any changes in the dimerization ability of TadA due to each mutation.

Consistent with our hypothesis and the entropy-based classification model, the D108N mutation leads to a modest 9.5% (ranging from 6.4% to 22.7%) increase in the A-to-I RNA-editing activity of ABE1.1 compared to ABE0.1. Moreover, the L84F mutation leads to a 22% (ranging from 18.5% to 28.7%), or almost 1.5-fold decrease, in the RNA-editing efficiency of the enzyme as compared to ABE0.1 across the six different RNA sites that were analyzed (Figure 4). These editing patterns were also observed at an additional UACG motif within RNA site 1, although the editing levels here were much lower than the other six sites (Supplementary Note 2).

**Figure 4:**
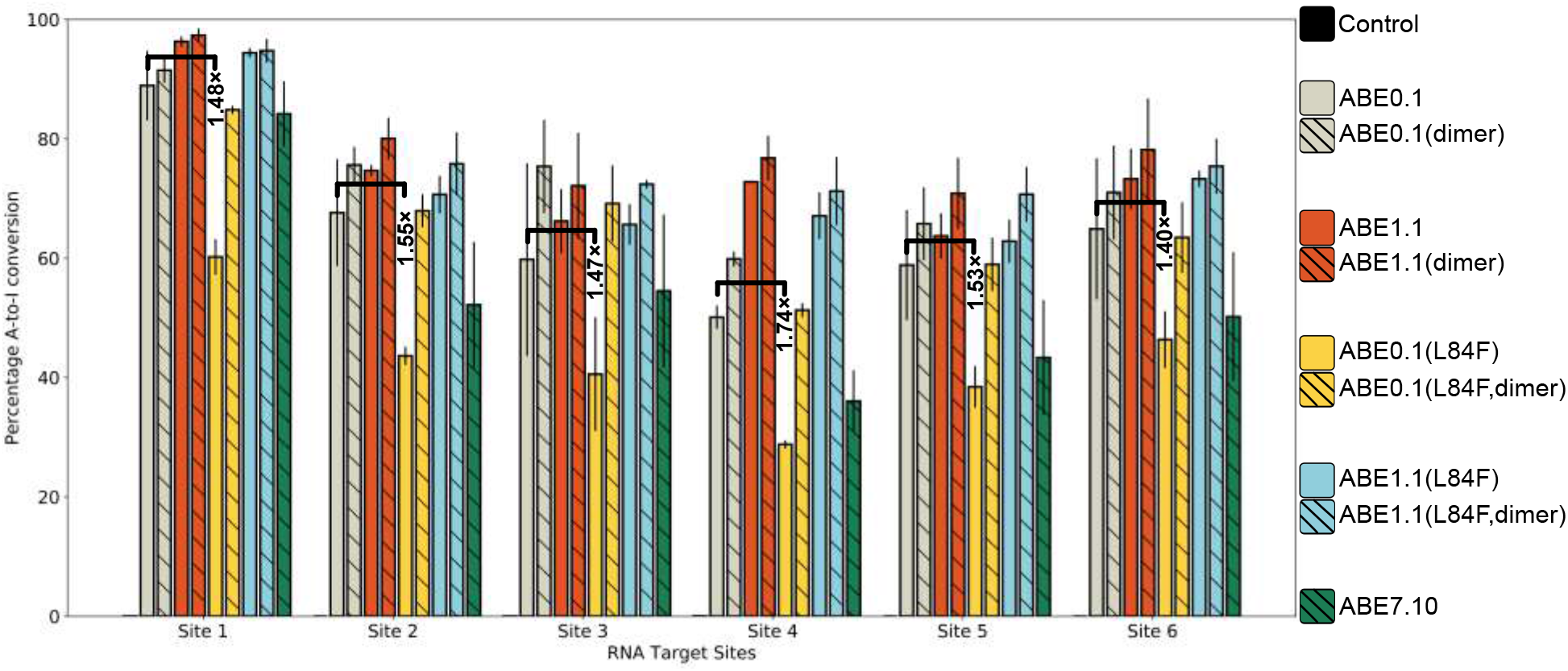
A-to-I base editing efficiencies in HEK293T cells by various ABE mutants at six different gRNA-independent RNA off-target sites. Fold-decrease values associated with the reduction in the RNA-editing upon incorporation of the L84F mutation in ABE0.1 are indicated. Values and error bars reflect the mean and SD of two independent biological replicates performed on different days.

This loss of function due to the L84F mutation can be restored either by dimerizing the protein with wtTadA (as ABE0.1(L84F, dimer)) or by adding the D108N mutation (as ABE1.1(L84F)). We speculate that in the case of ABE0.1(L84F, dimer) the observed increase in RNA-editing is due to the addition of the wtTadA subunit, which is capable of efficient RNA editing on its own (as in the ABE0.1 monomeric construct). In the case of ABE1.1(L84F), whose activity is comparable to ABE1.1, we observed a modest 7.3% (ranging from 3% to 16.9%) increase over ABE0.1. This restoration of the RNA-editing efficiency upon the combination of D108N and L84F mutations is especially interesting as it highlights the non-additive and epistatic effect that mutations can have on enzyme function. Thus, upon combining a high entropy (or high activity) mutation with a low entropy (or low activity) mutation, the resultant double-mutant exhibits high activity, rather than an average of the two activities. Furthermore, this double-mutant exhibits increased activity towards a different substrate (ssDNA).

Intriguingly, the RNA-editing activity of ABE7.10, which has 12 other mutations in addition to D108N and L84F, is slightly lower than that of ABE0.1 by 11.6% (ranging from 4.7% to 15.5%), or 1.2-fold (Figure 4). This observation further reinforces the non-additivity of the TadA* mutations identified using directed evolution of ABEs. The early mutations led to a broadening of the substrate specificity of TadA* (i.e. imparting upon TadA ssDNA-editing capabilities) and later mutations enhanced the ssDNA-editing activity while potentially suppressing the RNA-editing activity (as with the L84F mutation).

### Interaction and binding of RNA with TadA*

In Ref. 31, we demonstrated that the effects of individual mutations on the ssDNA-editing activity of ABEs can be studied using a minimalistic model of the system, comprised of the TadA* mutants and the nucleic acid substrates, while ignoring Cas9, which acts as a mere carrier of the nucleotide editing module to its target genomic locus. Moreover, it has been experimentally proven that the off-target RNA-editing by ABEs occurs in a Cas9 (or gRNA)-independent manner, which reinforces the notion that only the TadA* portion of the ABEs act on the RNA off-target substrates^10,13–16,32^

To understand the complex epistatic relationship between the L84F and the D108N mutations in the context of the RNA-editing activity of ABEs (Figure 3 and Figure 4), we modeled the ABE–RNA systems by combining the experimentally resolved structure of wild-type *E. coli* TadA (PDB :1Z3A^30^) and its native 14-mer RNA-hairpin substrate (5*t*-UUGACU**A**CGAUCAA-3’) (PDB :2B3J^33^). The RNA sequence in our simulation models, as well as the off-target RNA sites that we tested experimentally (Figure 4), have the same consensus sequence (-UACG-) as that reported previously^10,17^ (Figure S3). Moreover, the structures for these six RNA editing sites resemble the hairpin loop structure of the native target of TadA* that we simulated, further reinforcing the strong preference of TadA* for its native substrate (Figure S4). Having generated these models we then carried out MD simulations for each of the four TadA* mutants–RNA complexes for 1 microsecond and examined the trajectories for changes in interactions between individual TadA* residues and the nucleic acid substrate. Since the mutations we are interested in lie near the active site of the TadA*, we focus predominantly on the interactions between the nucleotide bases splayed in the active site, i.e., the target adenine and its 5’ and 3’ flanking bases (U**A**CG) and neighboring TadA* residues. To hone in on the amino acids in direct contact with these nucleotides, we carved out a 4 Å search radius around these bases, and then project the residues that lie within this sphere onto asteroid plots (Figure 5A to D). In the asteroid plots, the nucleotides in the active site are represented collectively as the central node, and the peripheral nodes correspond to all amino acids within the first interaction shell of the nucleotides in the active site. The size of the encircling nodes is proportional to the time that the corresponding residues spend within the first interaction shell of the RNA bases throughout the entire MD trajectory. The hydrogen bonds (H-bonds) between these residues and the RNA bases are depicted as arrows connecting the relevant nodes in each asteroid plot, with the thickness of the arrows being proportional to the stability of the H-bond itself, which is defined as the frequency of appearance of that H-bond during the simulation. The comparisons between the TadA*0.1/TadA*0.1(L84F), TadA*0.1/TadA*1.1, and TadA*1.1/TadA*1.1(L84F) mutants indicate that the D108N mutation leads to the formation of a favorable H-bond between the Asn108 residue and the U base flanking the target **A**. In fact, the TadA*1.1(L84F) mutant has the strongest interaction with RNA as the L84F mutation causes additional structural rearrangements surrounding the active site resulting in a double H-bond interaction with the RNA substrate through residue 152. The weak H-bond between D108 and the 2’-OH group of the flanking U base predicted by our simulations is also found in the crystallographic structure of the wtTadA-tRNA complex (PDB ID: 2B3J^33^). However, this weak H-bond does not appear in the TadA*0.1(L84F) mutant and is replaced by a much stronger H-bond in the TadA*1.1 mutant upon mutation of glutamate to asparagine at site 108. The formation of this stronger H-bond also leads to an increase in interactions between some of the peripheral residues (57, 59, 82, 85, 86, and 87) and the RNA bases in the active site, indicating a more stable conformation of the target adenine. A similar increase in the interaction induced by the H-bond formed by the D108N residue was also observed in our MD simulations of the TadA*–ssDNA complex.^31^ In the context of ssDNA-editing by ABEs, the D108N mutation in ABE1.1 leads to the onset of activity on DNA via the formation of this H-bond donation.^2,31^ However, in the context of RNA-editing efficiency, the D108N mutation, and consequent formation of the H-bond with RNA, only amounts to a slight increase in the activity due to wtTadA (ABE0.1) being already highly proficient in editing its native RNA substrate, as well as ssRNA in general (Figure 4).

**Figure 5:**
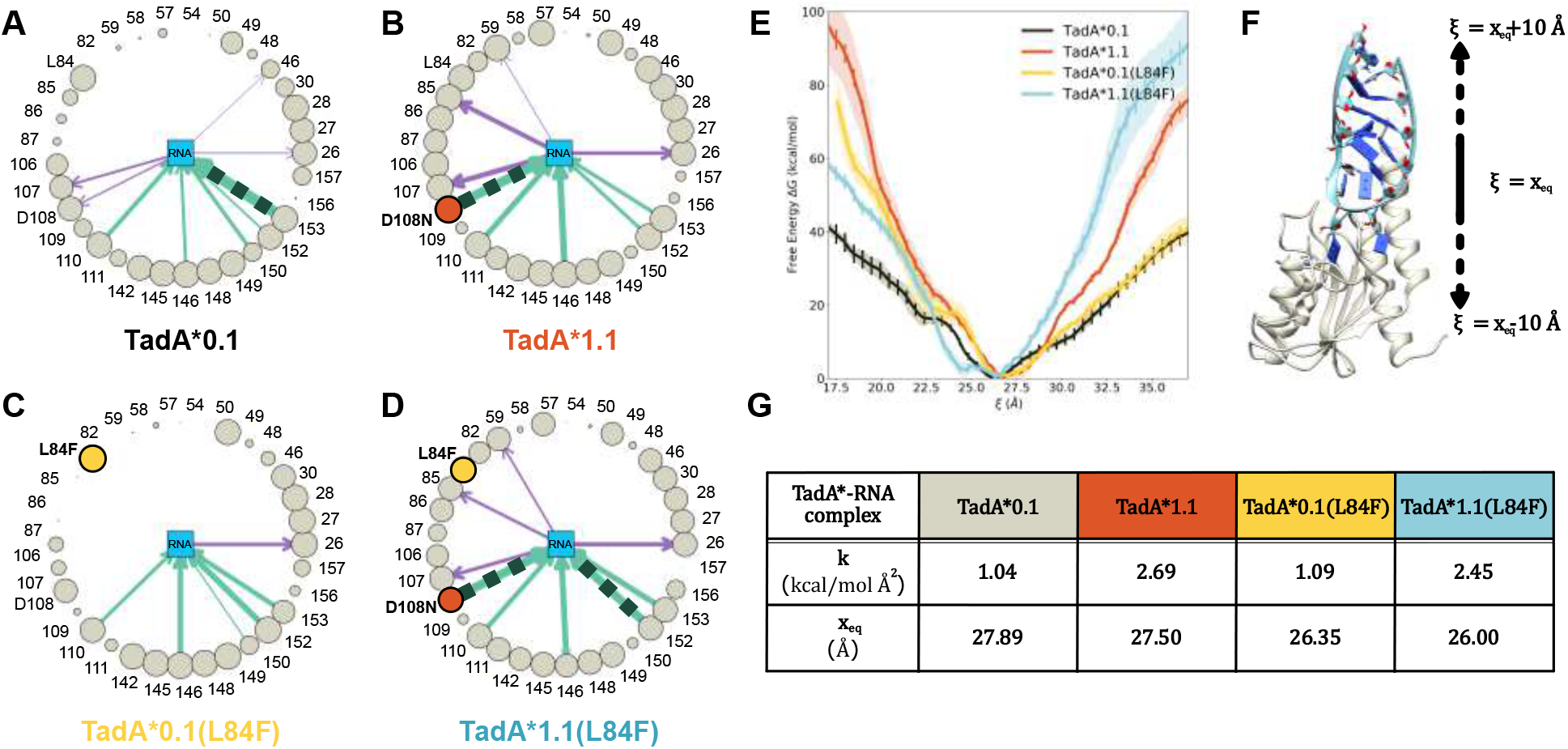
Asteroid plots for the analysis of the interaction of (A) TadA*0.1, (B) TadA*1.1, TadA*0.1(L84F), and (D) TadA*1.1(L84F) with substrate RNA. (E) Binding affinity comparisons for the various TadA*–RNA complexes. (F) The collective variable (*ξ*) used to monitor the binding/unbinding of the TadA*–RNA complexes. (G) Parameters associated to harmonic functions fitted to binding energy curves shown in (E).

Although the L84F mutation, unlike D108N mutation, is accompanied with a more pronounced effect on the RNA-editing activity of wtTadA (Figure 4), and is in fact a novel mutation in the first interaction shell of the RNA bases (Figure 3D), the comparison of the asteroid plots corresponding to TadA*0.1 and TadA*0.1(L84F) shows less drastic changes, than those observed in the TadA*1.1 asteroid plot. Specifically, the L84F mutation leads to an elimination of the weak H-bonds established by the 107 and 108 residues in TadA0.1. To quantify these differences, we performed umbrella sampling (US) simulations to determine the binding affinities of the various TadA*–RNA complexes. Starting from the equilibrated structure of each TadA*–RNA complex, we modeled the binding process using a collective variable (*ξ*) defined by the vector connecting the TadA* and RNA centers of mass, which was varied from 17 to 37 Å. We used this same collective variable to characterize the binding process in the analogous TadA*–DNA complexes in Ref. 31. The actual potential of mean force (PMF) associated with the RNA-binding process for each of the four TadA* mutants was calculated using the weighted histogram analysis method (WHAM).^34,35^ The work required to dissociate the TadA*1.1–RNA and TadA*1.1(L84F)–RNA complexes is larger than that required to dissociate the TadA*0.1–RNA complex. While these trends help explain the experimental RNA-editing efficiencies of these D108N mutants when compared to ABE0.1, they do not apply to L84F mutant for which we do not observe a decrease in the RNA-binding affinity relative between the TadA*0.1 and TadA*0.1(L84F) mutants. Although this observation may seem surprising, it, in fact, reciprocates the results of our previous study of the TadA*–ssDNA complex showing that mutations installed at later stages of the directed evolution process do not further enhance the binding affinity relative to TadA1.1, but instead most likely impact the catalytic activity of TadA*.^31^

As the subtle conformational changes that we observe between the asteroid plots of TadA*0.1 and TadA*0.1(L84F) (Figure 5 A and C) do not result in significant changes in the binding affinity between these two mutants, we thus sought to quantify the effects of these conformational changes on the catalysis.

### Water in the active site and implications for catalysis

The hydrodeamination reaction catalyzed by TadA involves a zinc-coordinated water molecule (hereafter referred to as the activated water molecule) that is deprotonated by the active site Glu^59^ residue (a highly conserved residue, see Figure 2) during the first step of the reaction. In addition to this activated water molecule, the active site also includes another structurally important water molecule (hereafter referred to as the bridging water) which acts as a bridge between Glu^59^ and the carbonyl backbone of Leu^84^ (Figure 6 A and B). Both water molecules are resolved in several high-resolution crystal structures of TadA homologs (Figure S5) which further reinforces their importance in the stabilization of the active site cavity.

**Figure 6:**
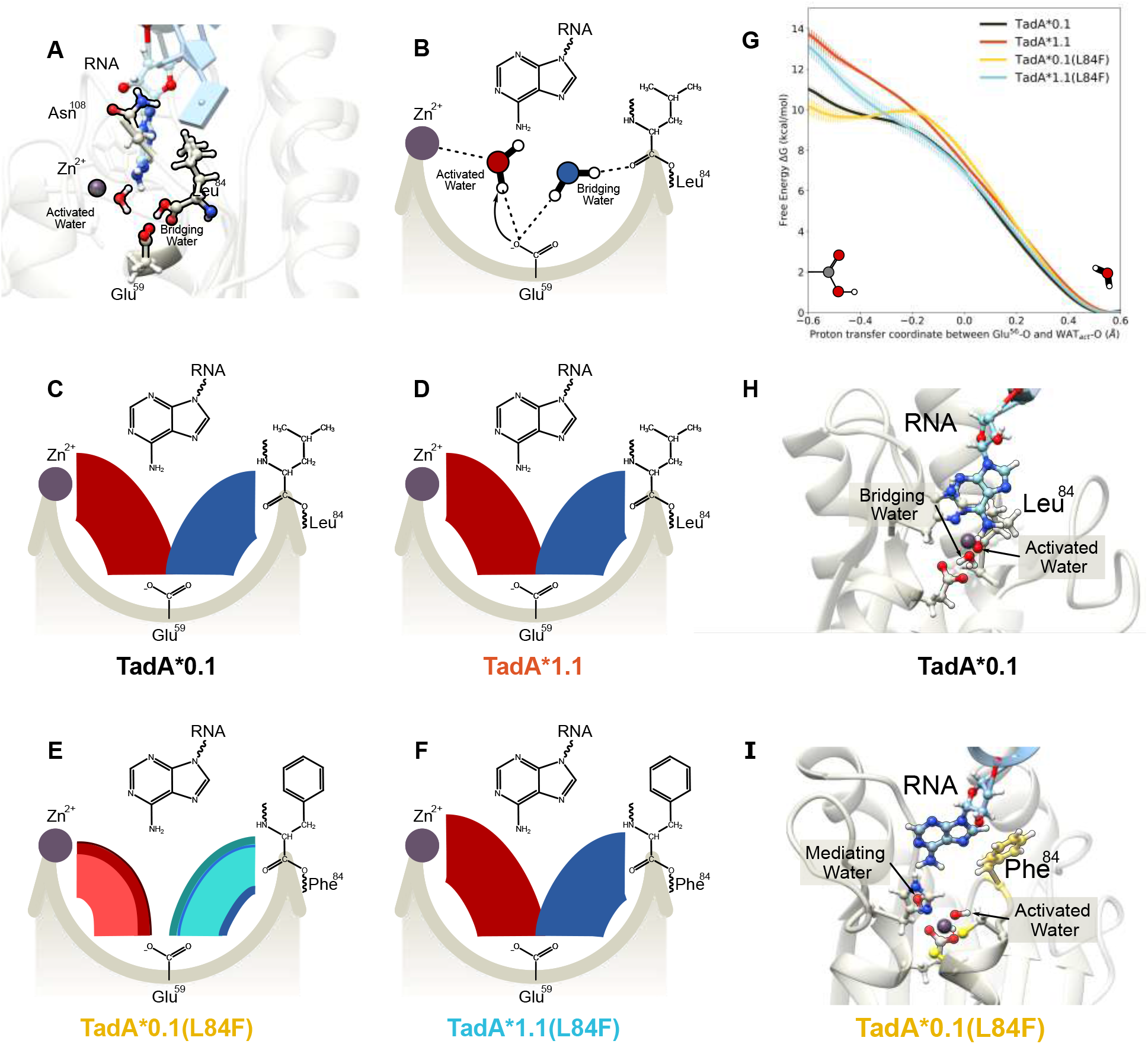
(A) Side view of the of the TadA*–RNA system highlighting the location of the catalytically relevant residues. The Zn^+2^ ion is coordinated by His^57^,Cys^87^, and Cys^90^ (not shown here for clarity) and a water molecule. This water molecule is activated by Glu^59^, which is also connected to another water molecule. This second water acts as a bridge between the Glu^59^ and the carbonyl backbone of residue 84. The target adenine is deep within the active site and residue 108 is farther away from the active site waters. (B) Simplified flat lay representation to highlight the interactions of active site waters. Modified chord diagrams to demonstrate the persistence of the active site waters for (C) TadA*0.1– RNA, (D) TadA*1.1–RNA, (E) TadA*0.1(L84F)–RNA, and (F) TadA*1.1(L84F)–RNA. (G) Reaction profile for the deprotonation of the activated water molecule the various TadA*– RNA systems. (H) Conformation of the TadA*0.1–RNA when the proton resides on Glu^59^. (I) Conformation of the TadA*0.1(L84F)–RNA when the proton resides on stability on the Glu59. The target **A**has moved back into the active site, towards the Phe^84^ and is separated from the active site residues by an additional water molecule - the mediating water.

To characterize the role played by these two water molecules in the active site of the various TadA*–RNA mutant systems, we analyzed the data from our MD simulations in the form of modified chord diagrams in (Figure 6 C, D, E, and F). The persistence of the activated water molecule is depicted in red in the left partitions while that of the bridging water is depicted in blue in the right partitions of the four panels. The thickness of each chord is proportional to the time spent by the corresponding water molecules in the active site (Figure S6). For the TadA*0.1–RNA and TadA*1.1–RNA systems, we found that these two water molecules are highly stable in their respective positions and do not undergo any diffusion throughout the entirety of our MD simulations (1 *μ*s). In contrast, for the TadA*0.1(L84F)– RNA system, both water molecules exhibit higher mobility and are exchanged several times with water molecules initially located in the bulk solution at the beginning of the simulation. We speculate that the hydrophobic-aromatic nature of the phenylalanine residue may be responsible for the decreased stability of both water molecules in the active site (Figure S7). The stability of the two water molecules is restored in the TadA*1.1(L84F)–RNA complex. This implies that the D108N mutation is capable of canceling out the destabilizing effects of the L84F mutation and effectively modulating the hydration of the active site, despite not engaging in any direct contact with either water molecules. We observe similar trends when comparing these mutations in the apo-TadA* simulations. Specifically, the apo-TadA*0.1(L84F) system shows an analogous increased flux of the two water molecules in the active site, which is again suppressed after the installation of the D108N mutation (Figure S8). Importantly, the changes in the persistence of these catalytically-relevant water molecules in the active site of the TadA*–RNA/TadA* systems (Figure 6 C, D, E and, F and S8) mirrors the changes in RNA-editing activity measured for the ABEs (Figure 4).

Since the first step of the adenine deamination reaction involves the deprotonation of the activated water molecule by the Glu^59^ residue, we speculate that the changes we observe in the stability of the active site water molecules may lead to changes in the reaction rates in the four TadA* mutants. Hence, for a more explicit comparison with the experimental catalytic data of these four TadA* mutants, we performed quantum mechanics/molecular mechanics (QM/MM) simulations to investigate this first step of the reaction mechanism. Owing to the high computational cost of simulating the entire system at QM level of accuracy, QM/MM simulations offer an ideal trade-off between accuracy and computational efficiency, by simulating the reaction centers with QM accuracy while the remaining system is treated at the MM level of theory. In our QM/MM simulations, the QM region encompasses the side chains of active site residues (His^57^, Glu^59^, Cys^87^, Cys^90^), Zn^+2^, and the activated water molecule, while all other atoms of the system are included in the MM region.^36^ All QM/MM simulations were initiated from configurations taken from the US window representing the PMF minima shown in Figure 5. In modeling the first step of the deamination reaction, configurations with undissociated activated water molecules define the reactant state, while configurations with the protonated Glu59 residue define the product state. In the transition state, the proton is equally shared by the activated water molecule and Glu^59^. To determine the energetics associated with this proton-transfer reaction, QM/MM umbrella sampling simulations were carried out along the proton-transfer coordinate, which is defined as the difference between the distances of the shared proton from the activated water and Glu^59^. The PMFs calculated using WHAM for all four TadA* mutants are shown in Figure 6G.

The PMF profiles indicate that only the TadA*0.1(L84F)–RNA complex is associated with a weakly stable product state (i.e. a protonated Glu^59^). At first glance, these results seem to be counter-intuitive and contrary to the experimental observation of a lower RNA-editing activity for the ABE0.1(L84F) mutant. However, upon further examination of the product state in the TadA*0.1(L84F)–RNA complex, we observed that proton transfer from the activated water to Glu^59^ is accompanied by the concomitant movement of the target adenine base away from the active site residue and towards the aromatic phenylalanine ring deeper into the active site forming in a staggered pi-stack with L84F residue (Table S1). The cavity formed as a result of this conformational change of the target adenine is filled by another water molecule (hereafter referred to as the mediating water) that may contribute to the following reaction steps, thereby altering the reaction mechanism for TadA*0.1(L84F). In the case of the TadA*0.1 (or TadA*1.1 and TadA*1.1(L84F)) system, our simulations predict that the active site retains its configuration, with the adenine base primed for the next steps of the reaction (Figure 6H). We thus hypothesize that the proximity of the adenine base prevents the transfer of the proton from the activated water molecule to Glu^59^. QM/MM umbrella sampling simulations carried out for the apo-TadA* mutants provide support for this hypothesis, showing the formation of a stable product state for all the TadA* mutants due to the lack of the adenine base (Figure S8G).

We thus conclude that the L84F mutation, a novel mutation at a low entropy residue site, affects the decrease in the activity of TadA* on the native RNA substrate through two key changes in its deamination chemistry. Firstly, this mutation destabilizes the two water molecules in the active site, which are both structurally and functionally relevant to the initiation of the deamination reaction. Secondly, it pulls the target adenine base away from the protonated Glu^59^, thereby making the subsequent reaction steps less feasible or leading to an alternate reaction pathway involving additional steps (e.g., through the mediating water). Our simulations indicate that the combination of the D108N mutation, which increases the RNA binding affinity, with the L84F mutation conserves the integrity of the active site by both stabilizing the two water molecules and positioning the target adenine appropriately for subsequent reaction steps, thereby, rescuing the catalytic activity of the ABE1.1(L84F) mutant (Figure 4).

## Conclusion

Through a systematic investigation of the various mutations that have been thus far identified in TadA*, our study re-traces the evolutionary trajectory followed by this enzyme using a data-driven approach that combined statistical models, MD simulations, and experimental assays.

The information contained in the naturally-evolved TadA homologs aids in rationalizing the effects of the mutations that have accumulated in the laboratory-engineered TadA* (Figure 2). We have demonstrated that mutations with a favorable impact on the RNA-editing activity of TadA* occur at residue sites having higher entropy, whereas mutations with an unfavorable impact on the RNA-editing activity occur at residue sites with lower entropy (Figure 3). Moreover, these low entropy sites when mutated to previously unvisited amino acids in the sequence space, such as the L84F mutation, can also have an adverse impact on the native function of the enzyme. Our experimental analyses also revealed that ABE0.1 has remarkably high gRNA-independent off-target RNA-editing and is even higher than the evolved ABE7.10 variant (Figure 4),^13,14,17^ a phenomenon which was previously overlooked.

These results indicate that such entropy-based scores, albeit being extracted from a highly RNA-biased dataset, can serve as a preliminary screen for site-directed mutagenesis and guide the library preparation for evolving future base editors with reduced off-target transcriptome editing activity. The most reliable inferences that can be derived from such biased datasets are related to the native RNA-editing functionality of the query sequence. Hence, we propose that this entropy-based tool be preferentially applied for the search of mutations that can suppress the inherent RNA-editing activity of potential base editors, a problem that, at present, cannot be solved using the traditional directed evolution methods.

Despite having reasonable success at predicting sites with low and high functionality for TadA*, the entropy-based tool has some inherent drawbacks. The assumption of using 0.5 as the dividing value for this classifier may be too simplistic. For future studies, instead of defining a single dividing value, we speculate that using a range of values (e.g., 0.5 ± 0.1) may lead to improvements in the predictive power of this classifier. Although the entropy-based model proves that laboratory-evolution plays by the rules set by natural evolution and that learning these rules from extant enzyme homologs can help guide future protein engineering endeavors, it is, by construction, not a generative model. That is, this model cannot be extended beyond the sequences in the dataset. It is limited by the diversity in the training dataset, which in our case is highly biased towards RNA-editing. Hence, we observed that directed evolution does explore regions of the sequence space that have not been explored by natural protein evolution (Table S2). Furthermore, statistical models for protein sequence analysis, like our entropy-based model, are known to perform poorly on predicting the relationship between co-evolving residues. ^37^ This is apparent in our mutual entropy analysis of 84 and 108 residue sites, where the entropy-based model is unable to decipher the correlations between these two sites (Supplementary Note 1) which can instead be understood through MD simulations of the TadA*–RNA complexes.

Using MD simulations we observed that L84F mutation decreases the experimental RNA-editing efficiency of TadA* by both destabilizing the active site water molecules and disrupting the conformation of the target **A**. These effects are fundamentally different from those induced by the D108N mutation which increases the experimental RNA-editing of TadA*. The D108N mutation increases the binding affinity between the substrate RNA and TadA* by establishing favorable H-bonds, a phenomenon that we have previously observed in our TadA*–ssDNA simulations.^31^ This additional binding affinity due to the D108N mutation alleviates the destabilizing effects of the L84F mutation, thereby restoring the experimental efficiency of TadA*1.1(L84F) mutant. Hence, the D108N and L84F mutations constitute an epistatic pair in the functional landscape of TadA*.^38^ The prevalence of such non-additive and complex relationships between the mutating residues make the task of rational protein design notoriously challenging, especially in cases such as that of TadA*, where the native RNA-editing fitness landscape is strongly coupled with the ssDNA-editing fitness landscape.

We anticipate that the synergistic combination of entropy-based models, experiments, and MD simulations described in this study can also enable researchers to apply similar multilevel strategies (informed by a combination of sequence, structure, and function of enzymes) to tackle the complex problem of the design of future base editors.

## Acknowledgement

The authors thank M. Norman for a Director’s Discretionary Allocation on the Comet GPU cluster at the San Diego Supercomputer Center. K.L.R. thanks C. Egan, E. Lambros, and S. Gu for helpful discussions.

## Author Contributions

The manuscript was written through contributions of all authors. K.L.R. performed all computer simulations. B.L.R. performed all experiments. K.R.G. and K.L.R. developed and analyzed the entropy-based model. A.C.K. and F.P. conceptualized and designed the research.

## Data and materials availability

HTS data have been deposited in the National Center for Biotechnology Information Sequence Read Archive database under accession code PRJNA687289. All other data needed to evaluate the conclusions in the paper are present in the paper and/or the Supplementary Materials. Additional data related to this paper may be requested from the authors.

## Supporting Information Available

The following files are available free of charge.

- Supplemental Figures (PDF)
- Detailed materials and methods (PDF)
- Details of force field parameters for the active site and input files for MD (ZIP file)

## Graphical TOC Entry

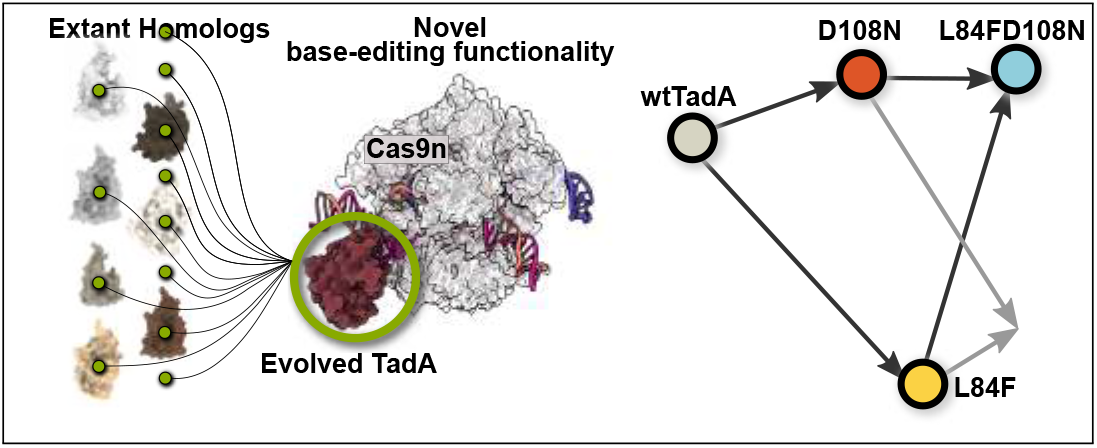

