## Supporting information for "Retracing the evolutionary trajectory of adenine base editors using theoretical approaches"

‖*Contributed equally to this work*

**3 Supplementary Figures 8**

**4 Supplementary Tables 23**

**5 Supplementary Sequences 26**

**6 Supplementary References 34**

### 1 Materials and Methods

#### 1.1 Data-driven statistical sequence analysis

##### 1.1.1 Data curation

Extant homologs were obtained using BLAST program[1] using *E. coli* wtTadA as the initial query sequence with an e-value cutoff of 0.1 in the SWISSPROT database.[2] This resulted in a dataset comprising of 75 homologs.

We then used special filters to reduce this dataset, by removing sequences with more than 40% gap percentage and to minimize redundant sequences with more than 95% identity to the query sequence. The final filtered dataset comprises of 35 homologs. Visual rationalization of the filtering process was performed with the help of dimensionality reduction through Principle Component Analysis (PCA), followed by K-means clustering. Given that each sequence in the aligned set of sequences contains 167 dimensions (length of query sequence is 167 amino acids), extracting the first two principle components from the PCA algorithm determines the 2-dimensional (2D) Cartesian coordinates of each of these sequences. Since the query of the multiple sequence alignment was *E. coli* wtTadA, it is important to realize that the coordinates of selected set of sequences are relative to the Cartesian coordinates of this sequence. We also calculated the distance of each sequence in 2D space from wtTadA. Additional phylogenetic analyses were performed in conjunction using ClustalW, and DENDROSCOPE[3] tool as well (Figure S1B).

To further substantiate the diversity of sequences selected by the filtering we implemented, a K-means clustering algorithm is run on the sequences upon the PCA dimensionality reduction. The optimal number of partitioning clusters (k) was determined to be four, through the conventional elbow method (Figure S1C).

##### 1.1.2 Entropy calculation

The resultant dataset was used to calculate the sequence entropy score, defined as follows:

$$S_i \ = \ -\sum_{i=1}^{N} p_i \, log_{20} \, p_i \quad \text{for} \quad i \in \{1, \ldots, L\} \quad \forall \quad \text{amino acids} \tag{1}$$

$$G_i \ = \ \frac{g_i}{N} \quad i \in \{1, \ldots, L\} \quad \forall \quad \text{gaps} \tag{2}$$

$$H_i \ = \ S_i \ + \ G_i \tag{3}$$

where $p_i$ refers to the statistical probability of having a particular amino acid at the site i and $g_i$ represented the total number of gaps at position i among all hits. The first term in the equation ($S_i$) represents the entropy value for amino acids at residue site i and the second term ($G_i$) represents the fraction of sequences with gaps (i.e. lack of any aligned amino acid after the MSA at that residue site). This treatment of the gap sites as a separate term ($G_i$) is done so as to retain the biochemical significance of presence of gaps, that such residue site are not functionally constrained. They can be (and have been) deleted from their parent sequences without damaging the activity of the enzyme.[4] The entropy scores obtained through Equation 3 were mapped on to the full-length structure of *E. coli* wtTadA that was generated by the missing C- and N-terminal residues using MODELLER[5] to PDB ID:1Z3A.[6]

##### 1.1.3 Binned Entropy Calculation

Binned entropy calculations are performed using a similar approach to that described in Equation 3. However, these entropies are not calculated wrt individual amino acids but rather a collections of similar amino acids (bins) defined on the basis of their chemical properties (Figure S2A). Thus, the binned entropy scores are evaluated as:

$$R_i = -\sum_{i=1}^{N} b_i \, log_6 \, b_i \quad \text{for} \quad i \in \{1, \ldots, L\} \quad \forall \quad \text{bins} \tag{4}$$

$$G_i = \frac{g_i}{N} \quad i \in \{1, \ldots, L\} \quad \forall \quad \text{gaps} \tag{5}$$

$$B_i = R_i + G_i \tag{6}$$

where $b_i$ refers to the statistical probability of a residue at site $i$ pertaining to bins shown in Figure S2A

#### 1.2 Experimental Methods

##### 1.2.1 Cloning

All plasmids used in this study were produced using USER cloning with ABE0.1[7] as a template, using Phusion U Hot Start DNA Polymerase (Thermo Fisher Scientific). All DNA vector amplification was carried out using NEB 10-beta competent cells (New England BioLabs). All plasmids were purified using the ZymoPURE II Plasmid Midiprep Kit (Zymo Research).

##### 1.2.2 Cell Culture

HEK293T (ATCC CRL-3126) cells were cultured in DMEM-GlutaMAX (Gibco) media supplemented with 10% (vol/vol) FBS (Gibco) and 100 U/mL penicillin-streptomycin (Gibco) at 37°C with 5% $CO_2$. Prior to transfection, the media was replaced with antibiotic-free media.

##### 1.2.3 Transfections

DNA samples for transfections were prepared with 750 ng of a base editor plasmid and 250 ng of a non-targeting sgRNA plasmid. Each sample was diluted to 12.5 $\mu$L with opti-MEM (Gibco). Lipofectamine 2000 (L2000, Invitrogen) was diluted with opti-MEM at a ratio of 1.5 $\mu$L L2000 to 11 $\mu$L optimem, with 12.5 $\mu$L of this mixture being added to each DNA sample. After 15 minutes of incubation at room temperature, each transfection sample was added to a well containing $9.5 \times 10^5$ HEK293T cells suspended in 250$\mu$L DMEM-GlutaMAX (Gibco) supplemented with 10% FBS.

##### 1.2.4 High-throughput RNA sequencing

Cells were lysed 36 hours after transfection with 300$\mu$l RNA lysis buffer (Zymo Research) and RNA was extracted with either a Zymo Quick-RNA Miniprep kit or Qiagen RNeasy mini kit following the manufacturers' instructions. RNA was reverse transcribed to produce cDNA using SuperScript III First-Strand Synthesis System (Invitrogen) following the manufacturer's instructions. Target sites were amplified from cDNA using two rounds of PCR. The first round used site-specific primers (Table S5) to amplify the target sequence from cDNA. Target amplification was confirmed with gel electrophoresis. The product from the first round PCR was used as a template in the second round of PCR, which added unique sets of p5/p7 Illumina barcodes to each sample. Amplification was confirmed with gel electrophoresis, then amplicons of similar size were pooled and gel purified on a 2% agarose gel. The target amplicon was excised from the gel and dissolved in 3 volumes QG buffer by incubating at 42°C for 10 minutes. After chilling on ice for 5 minutes, $\frac{1}{3}$ volume of 100% isopropanol was added and the sample was run through a Qiaquick PCR purification column on a vacuum manifold. The column was washed with 750$\mu$L of PE wash buffer and residual ethanol was removed by spinning the column at 16,000 g for 3 minutes. PCR products were eluted with 30$\mu$L HyClone water. Pooled libraries were quantified by Qubit and sequenced on an Illumina MiniSeq according to the manufacturer's protocol.

##### 1.2.5 HTS data analysis

Sequencing reads were demultiplexed in MiniSeq Reporter (Illumina) and individual FASTQ files were analyzed using a previously reported MATLAB script.[8]

#### 1.3 Molecular Dynamics Simulations

##### 1.3.1 System Setup

The TadA*0.1 model was built using the crystal structure of *E. coli* TadA (PDB ID: 1z3a).[6] Given the sequence homology between *S. aureus* TadA and *E. coli* TadA, we combined the *sa*TadA-RNA structure (PDB ID: 2B3J) with the TadA*0.1 model to build the TadA*0.1-RNA model.[9] The TadA*0.1 was transformed into the various ABE mutants using the `swapaa` command in Chimera.[10] For both apo-TadA* and TadA*-RNA models, all crystallographic water molecules within 3 Å distance of the surface of the protein or the RNA were preserved during the modeling procedure. All titratable residues were protonated using the H++ server using default settings.[11,12]

To parameterize the metal-containing active site of TadA* (comprising of $Zn^{+2}$, $His^{57}$, $Cys^{87}$, $Cys^{90}$, and an activated water molecule) we converted the two cysteines to their deprotonated state and used the MCPB.py approach at B3LYP/6-31G* level of theory.[13] The $Zn^{+2}$ ion and the side chains of its coordinating residues, along with the active site water were used to determine the bond and angle parameters between the $Zn^{+2}$ and the coordinating $S^{\gamma}$ atoms, $N^{\delta 1}$ atom, and the oxygen of the activated water molecule. To obtain the partial charges on the active site residues, a larger sub-system was chosen around the $Zn^{+2}$ ion. This larger sub-system was first optimized and then the optimized structure was used to obtain the partial charges through RESP fitting. The torsional terms involving the $Zn^{+2}$ ion were ignored for simplicity. The resultant model considered the active site as a fully-bonded metal center, with the $Zn^{+2}$ ion tetrahedrally linked to all its neighboring residues - including the activated water molecule.

To obtain a more realistic representation of the metal center and activated water molecule, we deleted the parameters associated with the $Zn^{+2}$ and the activated water and scaled the charge on the $Zn^{+2}$ ion appropriately. Hence, the ultimate model used in all the apo-TadA* and TadA*-RNA simulation involve a hybrid bonded- and non-bonded representation of the $Zn^{+2}$ active site, where the activated water molecule is no different from the bulk water molecules and can freely diffuse away from the ion. The partial charges and force field parameters are provided in the supplementary information folder.

The rest of the protein was represented using Amber ff14SB[14] and the RNA was represented using RNA.OL3 force field.[15–17] LEap tool from AmberTools was used to immerse the apo-TadA* and TadA*-RNA complexes into a pre-equilibrated truncated octahedron box of explicit TIP3P water, with a 15 Å buffer distance.[18] Varying number of $Na^{+}$ ions were added to each of the systems to maintain electroneutrality and the simulation cell was then replicated infinitely in three dimensions to impose periodic boundary conditions.

##### 1.3.2 Unbiased Molecular dynamics (MD) Simulations

To relieve any bad contacts that may have been present in the crystal structures or may have been introduced during the system preparation stages, all the apo-TadA* and TadA*-RNA models were first subjected to a multi-step energy minimization procedure using a combination of steepest descent and conjugate gradient algorithms. This minimization was carried out in four separate stages (((2000 steps of steepest descent + 3000 steps of conjugate gradient) × 4) total steps), each stage becoming less restrained than the previous one. During the first stage (5000 steps) only the solvent and the counter ions were allowed to move and all protein (or protein-RNA) atoms were restrained using a weight of 200 kcal/mol $Å^2$. During the second stage, all the heavy atoms of the protein (or protein-RNA) were restrained with a force of 200 kcal/mol $Å^2$, allowing only the hydrogen atoms and the solvent atoms to move freely. During the third stage, only the protein (or protein-RNA) backbone atoms were restrained with a force of 200 kcal/mol $Å^2$. During the fourth and final stage of minimization, all the restraints were removed and the entire system was allowed to relax freely.

The minimization was followed by heating to the temperature of 300 K, using a Langevin thermostat with the collision frequency of 2 $ps^{-1}$ to assign initial velocities to all the atoms in the system. This heating was followed by the equilibration of the systems in an isothermal-isobaric (NPT) ensemble with the Brendsen barostat for pressure scaling. Similar to the minimization procedure described above, the equilibration of the systems was also done in a multi-step manner (10 ns × 4), where restraints were gradually decreased (from 2.0 kcal/mol Å$^2$ to 1.0 kcal/mol Å$^2$ to 0.5 kcal/mol Å$^2$ and finally to no restraints) from all the protein (or protein-RNA) atoms . Long-range electrostatics were evaluated using the Ewald method with the non-bonded energy cutoff at 10 Å.[19] The SHAKE algorithm was used for constraining all bond distances involing hydrogen atoms.[20]

These equilibrated structures were used to initiate the 1 $\mu$s simulations as well as the biased MD simulations used to calculate the binding affinity between the various protein-RNA complexes. All simulations were propagated in time using the velocity Verlet algorithm with a time step of 2 fs. All simulations were conducted using the CUDA accelerated version of PMEMD Amber18.[21–24]

###### 1.3.3 Binding energy Simulations

Starting from the equilibrated structures obtained from the unbiased MD, as described above, we calculated the binding energy profiles for the TadA*-RNA complexes. This was done in two parts : first, a preliminary estimation was generated using steered MD (SMD) simulations,[25] which was followed by confirmatory umbrella sampling (US) calculations.[26]

The collective variable ($\xi$) used to monitor the binding or unbinding process was defined as the distance between the centers of mass (COM) of the TadA* and the RNA ( Figure 5F). Using a constant rate of 0.1 Å/ns pulling (and pushing) the TadA*-RNA complexes were dissociated (and associated) beyond their equilibrated distance ($\approx$ 17 Å) by ± 10 Å.

Configurations from these SMD binding profiles were used as the seeds for longer US simulations. The coordinate space ($\xi \in$ [17, 37] Å; step 0.5 Å) was divided to generate a set of 41 discrete US windows. Each US window was subjected to a production stage for 5 ns under the umbrella restraints with a force constant, k, of 40 kcal/mol Å$^2$.

Four independent US simulations were carried out for each of the 41 windows for all the TadA*-RNA complexes, which were then used to determine the potentials of mean force (PMFs) representing the free energy profiles associated with the binding process along the $\xi$.

Thus, a total of 820 ns (20 ns × 41 windows) was sampled for each TadA*-RNA complex and this led to the accumulation of 100000 instantaneous values scanning along the $\xi$ coordinate. Due to the external bias applied on the system through harmonic restraining , this biased probability distribution (Figure S9) is uniformly distributed along $\xi$. To reveal the true unbiased probability distribution and thus, the PMF along $\xi$, we made use of the WHAM algorithm,[27,28] with a convergence threshold of $10^{-8}$.

With 4 independent PMF profiles for each of the TadA*-RNA complexes, the error in convergence due to the sampling was calculated as a the standard deviation of the 4 datasets. Additional error analysis was performed using the block averaging method to evaluate the uncertainty associated with the normalization procedure implemented within the WHAM algorithm.[29]

###### 1.3.4 QM/MM Simulations

Through a hybrid quantum mechanical/molecular mechanical (QM/MM) approach the free energy changes for the deprotonation of the activated water molecule by the Glu$^{59}$ residue for the TadA* (and TadA*-RNA) models were computed for the various mutants.[30]

As this first step involves the transfer of a proton from the water (activated by $Zn^{+2}$) to the Glu$^{59}$, the QM subsystem consisted of the side chains of the active site residues (His$^{57}$, Glu$^{59}$, Cys$^{87}$, and Cys$^{90}$), the $Zn^{+2}$ ion, and the activated water for both the apo-TadA* and TadA*-RNA models. These QM atoms were treated using self-consistent charge density functional tight binding (SCC-DFTB) method implemented within Amber18, which has been shown to have good accuracy at nominal computational expense despite being a semiempirical model.[31,32] The atoms beyond this active site cluster were represented the MM subsystem and were treated using the force fields as in the unbiased MD.

The QM subsystem was electrostatically embedded into the MM subsystem, and the bonds spanning the two subsystems were capped using hydrogens as the link atoms. The net charge on the QM region was -1. The electrostatics of the QM region were cutoff beyond 9 Å distance. No SHAKE was implemented in the QM region as the reaction to be simulated involves a proton and the time step was reduced to 0.5 fs. All QM/MM simulations were conducted with the QM(DFTB)/MM implementation in the sander module of Amber18.

For the apo-TadA* systems we started from the equilibrated conformations extracted from the unbiased MD and for the TadA*-RNA systems we started from the conformations extracted from the minima in the binding energy profile, to initiate the catalytic deprotonation reaction. These structures were further equilibrated using the QM/MM scheme for 500 ps, which was followed by a QM/MM SMD simulation to explore the reaction profile of the deprotonation.

The difference of the distances between the Wat $O$ and shared proton and the $\mathrm{Glu}^{59}\,O$ and shared proton, was chosen as the collective variable ($\xi$) to monitor the deprotonation. Using the QM/MM SMD scheme with a constant pulling speed of 0.0024 Å/ps and a force constant of 200 kcal/mol Å$^2$, the shared proton was forced away from the water and onto the $\mathrm{Glu}^{59}\,O$.

From this preliminary estimate of the reaction profile of the system, snapshots were extracted at every 0.1 Å, from $\xi \in$ [-0.6, 0.6] Å, to seed 13 discrete windows, to conduct 1950 ps (13 windows $\times$ 150 ps) of simulations under umbrella restraints. This led to the accumulation of 300000 instantaneous values of the $\xi$ for each individual window. To obtain the unbiased probability distribution from this raw data, and thus, the PMF along $\xi$, we again made use of the WHAM algorithm with a convergence threshold of $10^{-8}$ (SI Figure Figure S10 and Figure S11).

With 3 different PMF profiles for each of the TadA* and TadA*-RNA models, the error in the average reaction profile was estimated as the standard deviation of these sets.

#### 1.4 Analysis Protocol

The trajectories from the simulations of the various apo-TadA* and TadA*-RNA systems were analyzed using the cpptraj tool.[33,34] For creating the asteroid plots, we first identified the amino acid residues within the primary interaction shell around the nucleotides in the acitve site (`-UACG-`) and then using these residues in the cpptraj distance-based `mask` we analyzed the trajectories of the unbiased MD. The atom-list per frame data obtained using the cpptraj module was re-normalized to give the percentage residue contacts, using the following formula:

$$\text{Percentage contact} = \frac{\text{Total atomic contact during all frames} \times 100}{\text{Number of atoms in the amino acid} \times \text{Total number of frames}} \tag{7}$$

The average number of hydrogen-bonding interactions between the residues in the primary interaction shell and the RNA bases was computed using the `hbond` feature in cpptraj with the default hydrogen-bond definition (3 Å distance between donor and acceptor atoms, and 135° angle between the donor, hydrogen, and acceptor atoms) (Figure S12).

To generate the modified chord diagrams, we tracked individual waters using a similar distance-based `mask` but this time restricted to only water oxygens and ignoring all the other atoms of the systems (SI Figure S6). The visualization of all the trajectories was rendered using Chimera,[10] the graphical data was plotted using Matplotlib,[35] and the curve-fittings (SI Figure S9) were done using.[36]

### 2 Supplementary Notes

#### 2.1 Supplementary Note 1

To understand this non-additive behavior in ABE1.1(L84F) we again rely on our entropy-based analysis. As residues 84 and 108 are in direct contact (see the three-dimensional structure of TadA* in (Figure 3F)), we speculate that each of these residues may encode useful information regarding the nature of the other. To test this hypothesis, we define the covariance of two residue sites $i$ and $j$ using mutual information as:

$$\text{Mutual Information} \; = \; \text{MI}_{i,j} \; = \; H_i + H_j - H_{i,j} \tag{8}$$

where $H_i$ and $H_j$ are the self-entropy scores calculated using Equation 1, and $H_{i,j}$ is the joint entropy for the occurrence of a certain pair of amino acids at sites $i$ and $j$. This computation leads to a mutual information score between residues 84 and 108, $MI_{84,108}$, equal to 0.245. In the context of all the $MI$ values calculated for the entire sequence of TadA, $MI_{84,108}$ is in the 74th percentile (Figure S13), indicating a somewhat weak correlation between the two sites. However, it should be noted that in our dataset site 84 is never encountered to be a phenylalanine (Figure 3D). Thus, there is no combination of 84F and 108N in our dataset. This indicates that mutual information is not the appropriate metric to understand the nature of the interaction between residues 84 and 108.

#### 2.2 Supplementary Note 2

An additional `-UACG-` motif was identified within the RNA editing Site 1 amplicon (referred to as Site $1'$ henceforth) which was previously unreported.[37] Editing levels at this site are much lower than the other six sites, but otherwise follow the same patterns when treated with the different ABE variants (Figure S14. Interestingly, there is a drastic difference in editing levels at this site between ABE0.1 and ABE1.1. Introduction of the L84F mutation to ABE1.1 drastically lowers editing levels, and even lower editing is seen with ABE7.10. The secondary structure for Site $1'$ is highlighted in Figure S3.

### 3 Supplementary Figures

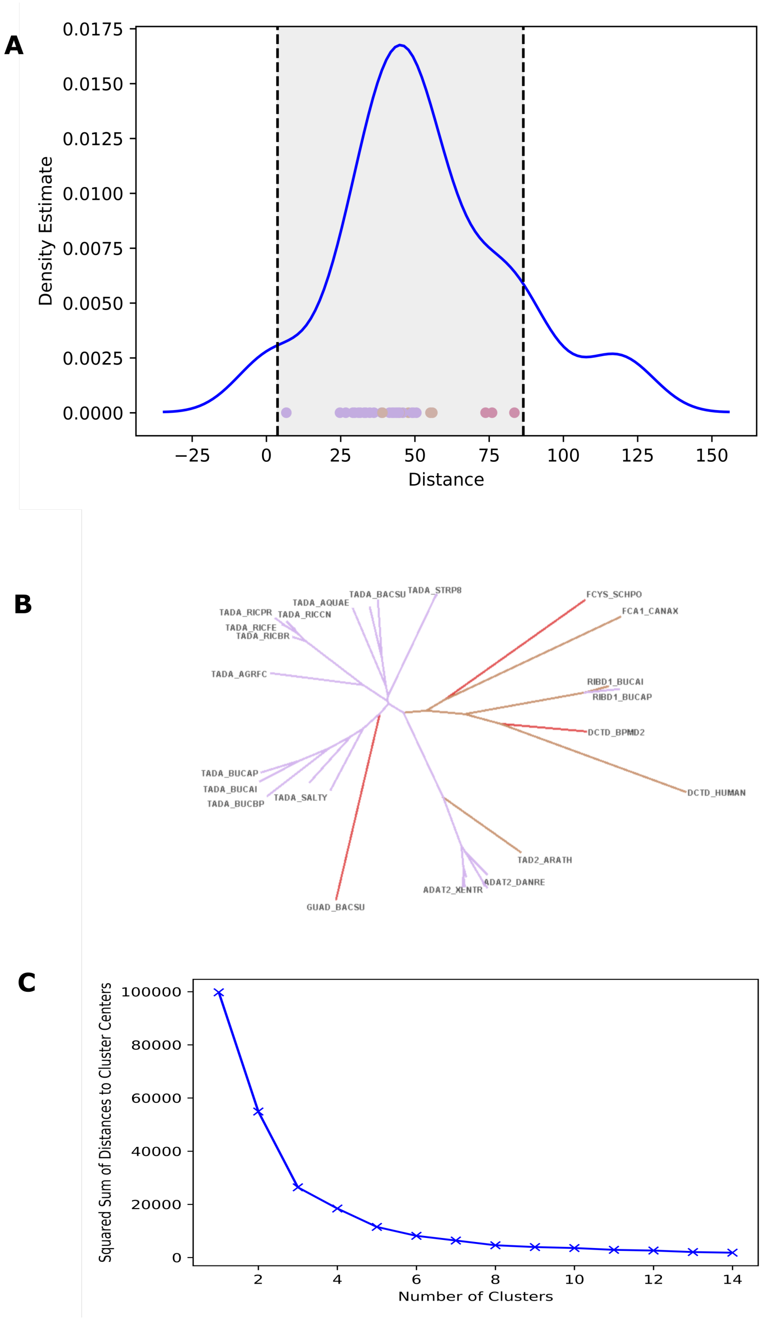

Figure S1: (A) Kernel density estimate (KDE) showing the distance of sequences from *E. coli* wtTadA from the first 2 principal components. (B) Phylogenetic analysis of the alignment for *E. coli* wtTadA. (C) Elbow plot to determine the optimal number of clusters to be used in k-means clustering.

**A**

| Amino Acid | Category |
|---|---|
| Arginine, Histidine, Lysine | Positively charged |
| Aspartic Acid, Glutamic Acid | Negatively charged |
| Serine, Threonine, Aspargine, Glutamine | Polar uncharged |
| Alanine, Valine, Leucine, Isoleucine, Methionine | Hydrophobic-aliphatic |
| Phenylalanine, Tyrosine, Tryptophan | Hydrophobic-aromatic |
| Cysteine, Selenocysteine, Glycine, Proline | Special |

**B**

**C**

**D**

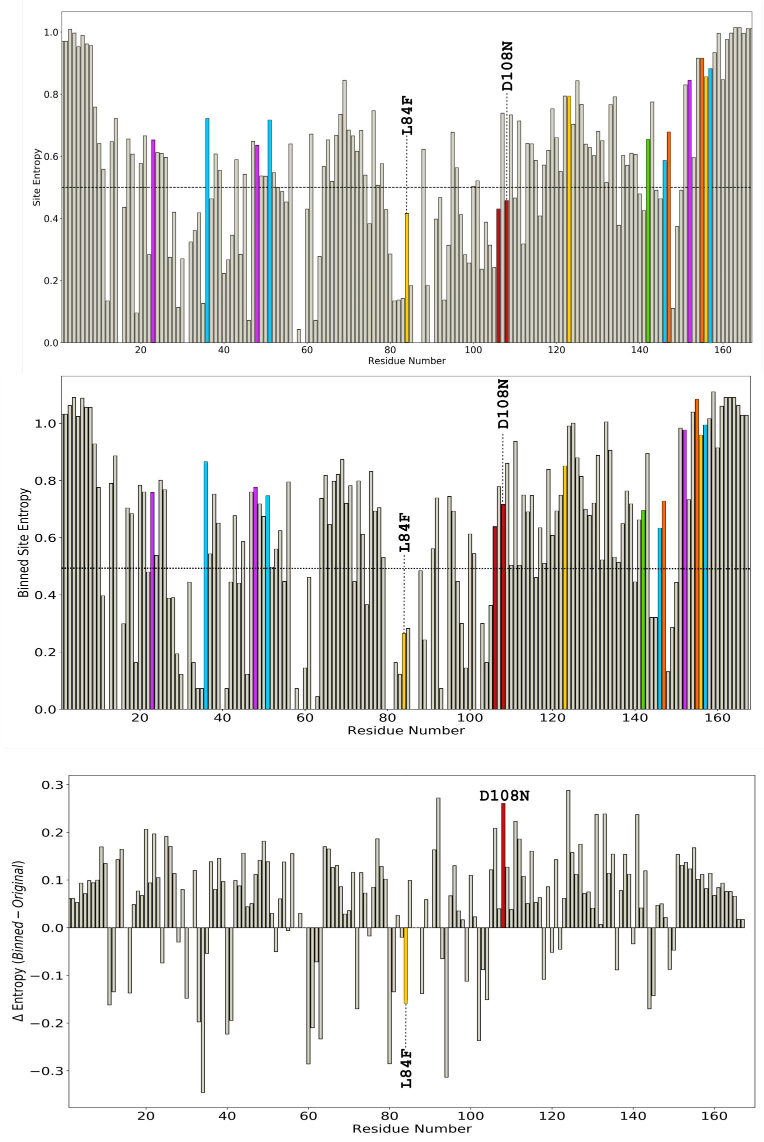

Figure S2: (A) Categories into which each of the 20 amino acid is placed for analyzing site-specific amino acid diversity. (B) Entropy-based analysis of ABE7.10 mutations. (C) Binned-entropy based analysis of ABE7.10 mutations. (D) Difference in the binned and regular entropy values to highlight the increase in entropy of site 108 and decrease in the entropy of site 84.

**Experimentally tested sites**

RNA site 1 (DNAJB1) (chr19: 14518195)

GCGCTACCACCCGGACAAGAACAAGGAGCCCGGCGCCGAGGAGAAGTTCAAGGAGATCGCTGAGGCC**TACG**ACGT

GCTCAGCGACCCGCGCAAGCGCGAGATCTTCGACCGCTACGGGGAGGAAGGCCTAAAGGGGAGTGGCCCCAGTGG

CGGTAGCGGCGGTGGTGCCAATGGTACCTCTTTCAGCTACACATTCCATGGAGACCCTCATGCCATG

RNA site 2 (MTA2) (chr11: 62594034)

GGGATTCGTTCAAGCTCACAGCCAGCAGCCAAGCGTCAGAAACTAAACCCAGCTGATGCCCCCAATCCTGTGGTG

TTTGTGGCCACAAAGGATACCAGGGCCC**TACG**GAAGGCTCTGACCCATCTGGAAATGCGGCGAGCTGCTCGCCGA

CCCAACTTGCCCCTGAAGGTGAAGCCAACGCTGATTGCAGTGCGGCCCCCTGTCCCTCTACCTGCACCCTCACATC

RNA site 3 (PTBP2) (chr1: 96813053)

AGATTTTGGTAATTCCCCATTGCATCGTTTTAAGAAACCTGGATCCAAAAATTTTCAAAACATTTTTCCTCCTTCT

GCCACCCTTCACCTATCTAATATCCCTCCATCAGTAGCAGAAGAGGATC**TACG**AACACTGTTCGCTAACACTGGGG

GCACTGTGAAAGCATTTAAGTTTTTTCAAAGAGATCACAAAATGGCTCTTCTTCAGATGGCAACAGTGGAAGAAGC

TATTCAGG

RNA site 4 (SAP30BP) (chr17: 75703316)

CAGAACCCCCTGGCAGATGTTCAAATCACTTGCAAGACAAGATCCAGAAGCTTTATGAACGAAAGATAAAGGAGGGA

ATGGATATGAACTACATTATCCAAAGGAAGAAAGAATTTCGGAACCCTAGCATC**TACG**AGAAGCTGATCAGTTCTGT

GCCATTGACGAGCTTGGCACCAACTACCCAAAGGATATGTTTGATCCCCATGGCTGGTCTGAGGA

RNA site 5 (LCMT1) (chr16: 25164711)

ATTGCCAACACTCCTGATAGCTGAATGTGTGCTGGTTTACATGACTCCAGAGCAGTCCGCAAACCCCTGAAGTGGGC

AGCCAACAGTTTTGAGAGAGCCATGTTCATAAAC**TACG**AACAGGTGAACATGGGTGATCGGTTTGGGCAGATCATGA

TTGAAAACCTGCGGAGACGCCAGTGTGACCTGGCGGGAGTGGAGACCTGCAAGTC

RNA site 6 (SCAP) (chr3: 47420696)

CTCCATCCGGCGAATGTCAATGGCTAGCAGACCTGAACAAGCGACTGCCCCCTGAGGCCTGCCTGCCCTCAGCCAAG

CCAGTGGGACAGCCAACGCGC**TACG**AGCGGCAGCTGGCTGTGAGGCCGTCCACACCCCACACCATCACGTTGCAGCC

GTCTTCCTTCCGAAACCTGCGGCTCCCCAAGAGGCTGCGTGTTGTCTACTTC

**Sequence used in the simulation**

UUGAC**UACG**AUCAA

Figure S3: RNA sequences tested in the experimental analyses and simulation models. The amplicon sequences pertain to the cDNA used for sequencing analysis. Thus, every T should be considered a U in RNA. The consensus sequence for all sites tested is hence -UACG-, which is the same as the native RNA substrate sequence used in the simulation models.

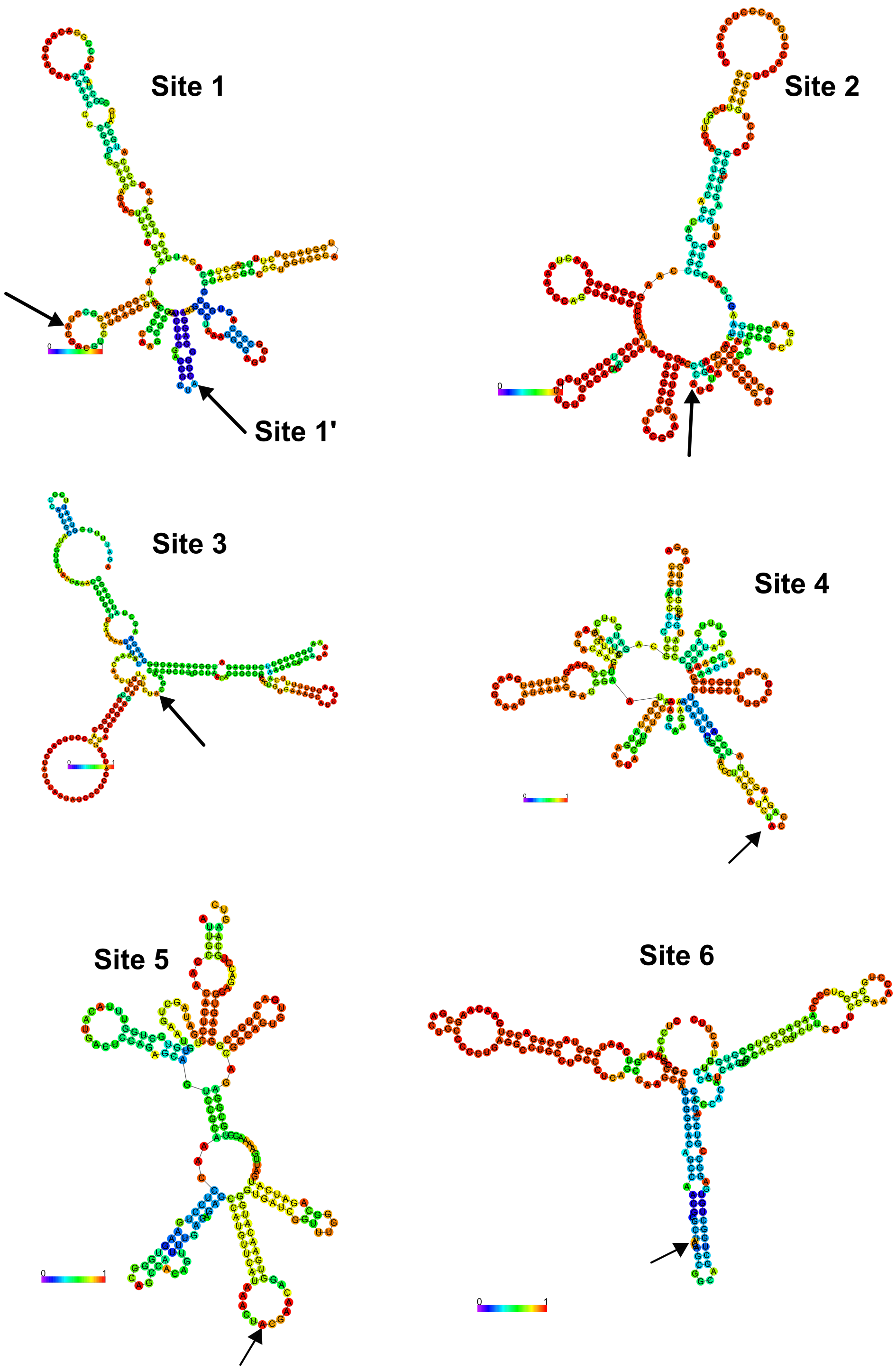

Figure S4: Secondary structures of the six RNA sites that are tested experimentally. The structure were determined using the minimum free energy method of the RNAfold webserver with the default parameters.[38] The arrows here indicate the Adenine base that is being edited by the various ABEs. The bases are colored by their base-pairing probabilities. For unpaired regions the color denotes the probability of being unpaired. All sites, expect site 6, predict the target adenine to occur as unpaired hairpin loop regions.

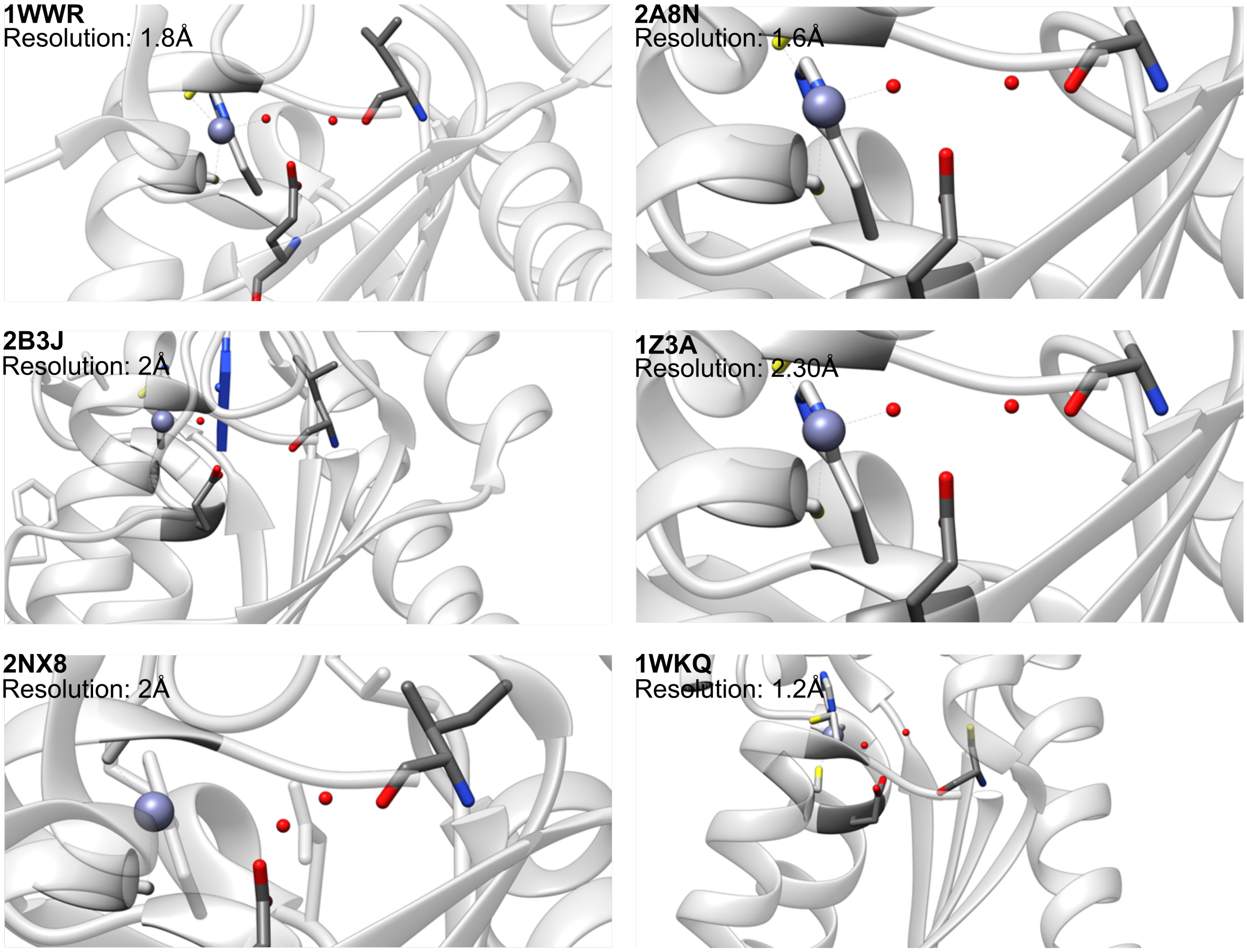

Figure S5: Crystal structures of *ec*TadA homolog highlighting the active site architecture and the relative position of the two critical waters, the activated water and the bridging water, hydrogen bonded to the same glutamate oxygen atom.

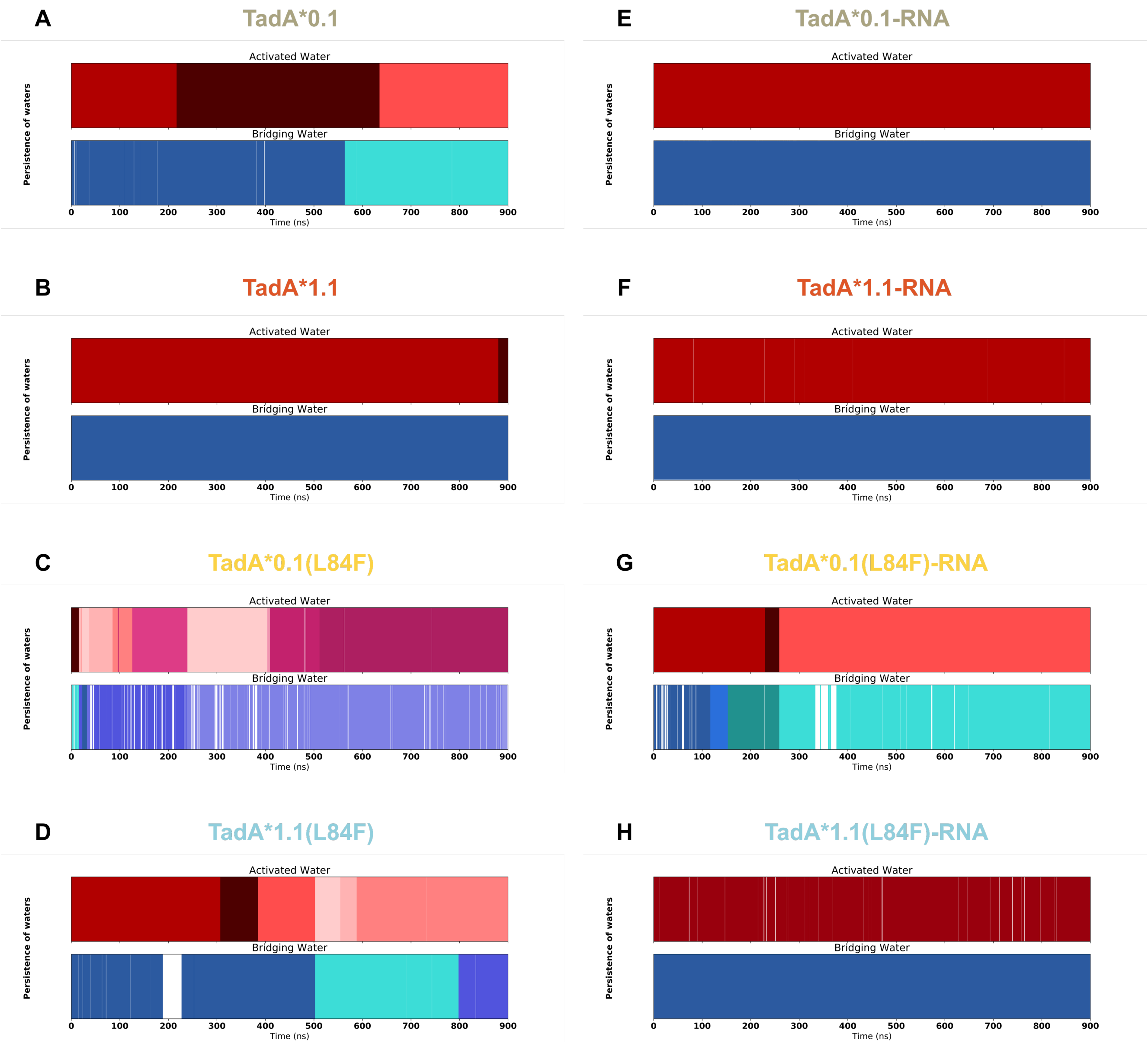

Figure S6: Persistence of the activated and bridging water at their respective positions calculated through the analysis of the unbaised MD trajectories of the apo-TadA* models ((A) to (D)) and for TadA*-RNA models ((E) to (G)). The different colors signify a unique water molecule that visited these critical water positions. Note that the data was obtained using a distance-based `mask` of 3.5 Å from the $Zn^{+2}$ ion, for activated water and from the peptide backbone of the 84 residue, for the bridging water.

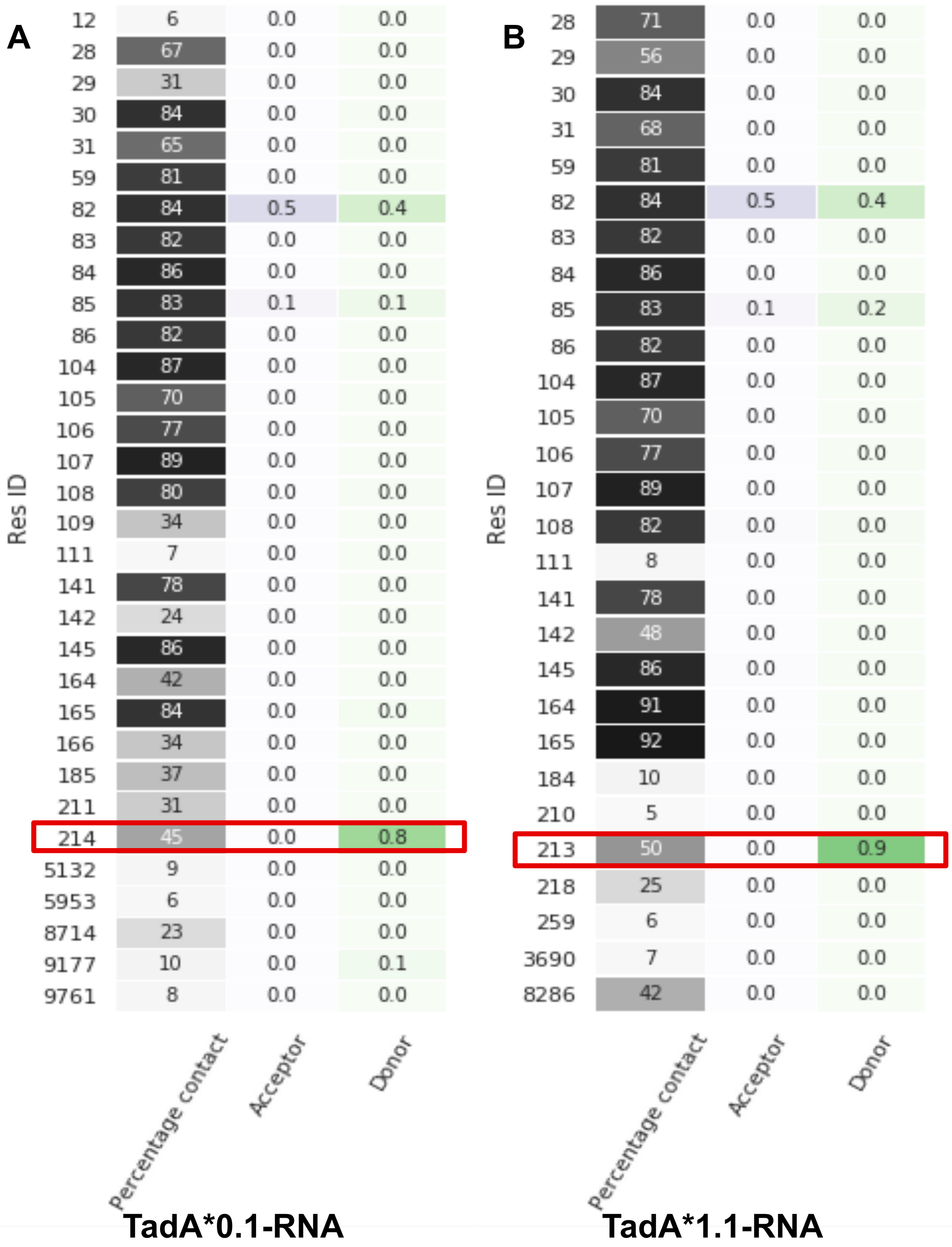
A
Res ID
12 6 0.0 0.0
28 67 0.0 0.0
29 31 0.0 0.0
30 84 0.0 0.0
31 65 0.0 0.0
59 81 0.0 0.0
82 84 0.5 0.4
83 82 0.0 0.0
84 86 0.0 0.0
85 83 0.1 0.1
86 82 0.0 0.0
104 87 0.0 0.0
105 70 0.0 0.0
106 77 0.0 0.0
107 89 0.0 0.0
108 80 0.0 0.0
109 34 0.0 0.0
111 7 0.0 0.0
141 78 0.0 0.0
142 24 0.0 0.0
145 86 0.0 0.0
164 42 0.0 0.0
165 84 0.0 0.0
166 34 0.0 0.0
185 37 0.0 0.0
211 31 0.0 0.0
214 45 0.0 0.8
5132 9 0.0 0.0
5953 6 0.0 0.0
8714 23 0.0 0.0
9177 10 0.0 0.1
9761 8 0.0 0.0
Percentage contact
Acceptor
Donor
TadA*0.1-RNA
B
Res ID
28 71 0.0 0.0
29 56 0.0 0.0
30 84 0.0 0.0
31 68 0.0 0.0
59 81 0.0 0.0
82 84 0.5 0.4
83 82 0.0 0.0
84 86 0.0 0.0
85 83 0.1 0.2
86 82 0.0 0.0
104 87 0.0 0.0
105 70 0.0 0.0
106 77 0.0 0.0
107 89 0.0 0.0
108 82 0.0 0.0
111 8 0.0 0.0
141 78 0.0 0.0
142 48 0.0 0.0
145 86 0.0 0.0
164 91 0.0 0.0
165 92 0.0 0.0
184 10 0.0 0.0
210 5 0.0 0.0
213 50 0.0 0.9
218 25 0.0 0.0
259 6 0.0 0.0
3690 7 0.0 0.0
8286 42 0.0 0.0
Percentage contact
Acceptor
Donor
TadA*1.1-RNA

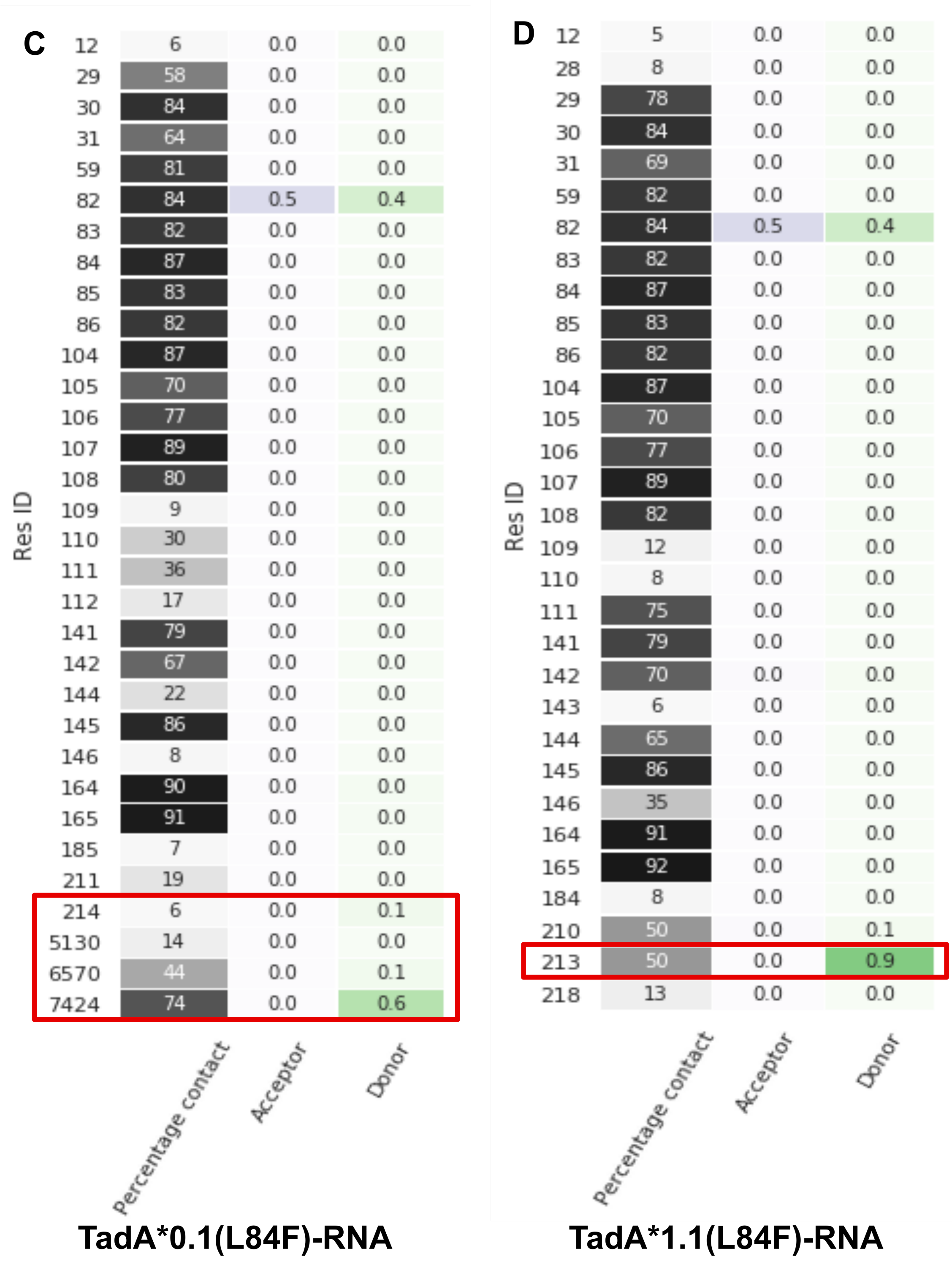

Figure S7: Percentage contact and the fractional H-bonding of the 84 residue. For (A) TadA*0.1, (B) TadA*1.1, (C) TadA*0.1(L84F) and (D) TadA*1.1(L84F) in complex with RNA.The intensity of the colors in the columns signifies the magnitude of the percentage contact and the H-bonding strength. The waters bridging (i.e. the bridging waters) the backbone of 84 residue and the Glu[59] residue are highlighted for each mutant.

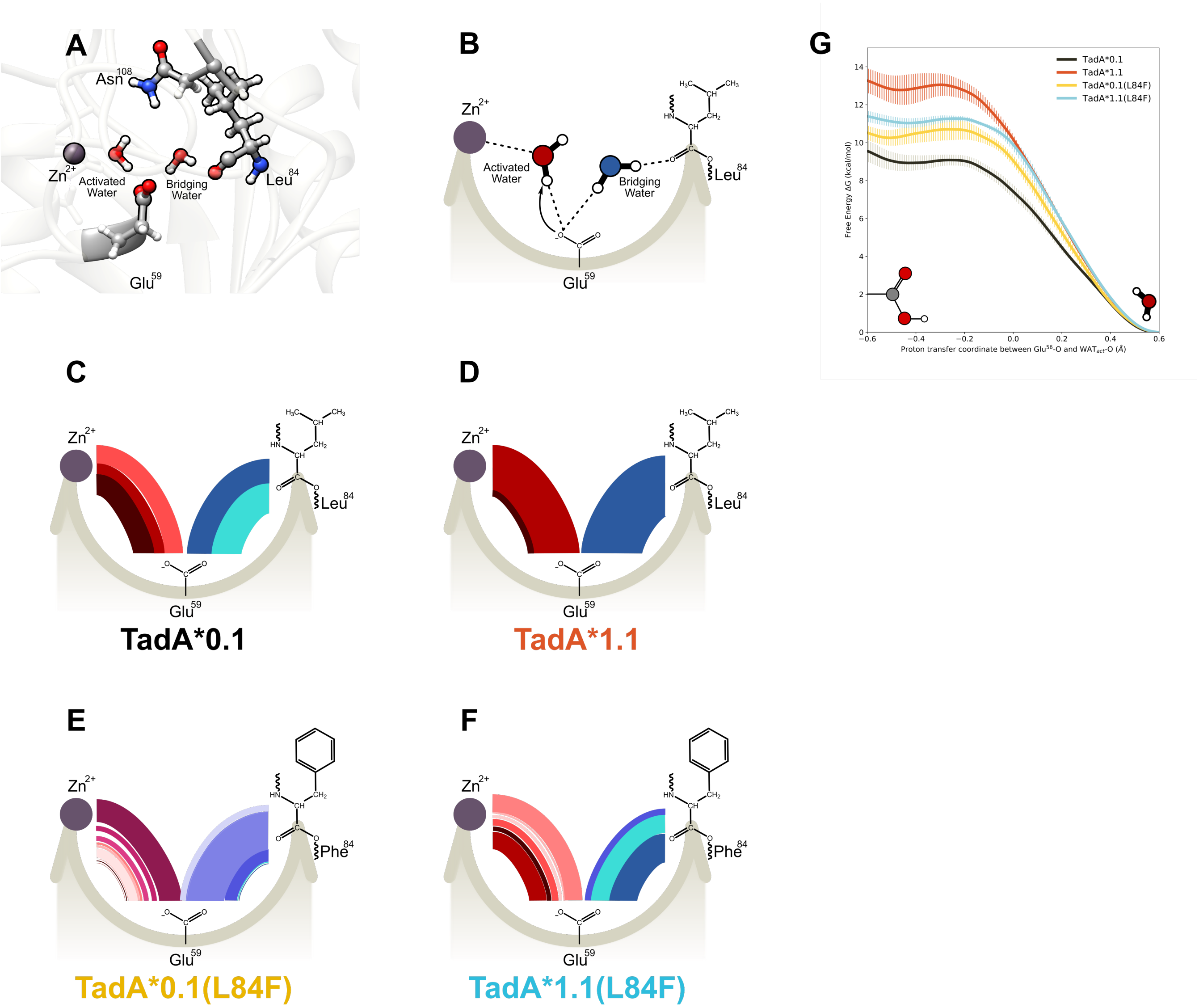

Figure S8: (A) Side view of the of the apo-TadA* models highlighting the location of the catalytically relevant residues. The $Zn^{+2}$ion is coordinated by $His^{57}$,$Cys^{87}$, and $Cys^{90}$ (not shown here for clarity) and a water molecule. This water molecule is activated by $Glu^{59}$,which is also connected to another water molecule. This second water acts as a bridge between the $Glu^{59}$ and the carbonyl backbone of residue 84. The target adenine is deep within the active site and residue 108 is farther away from the active site waters. (B) Simplified flat lay representation to highlight the interactions of active site waters. Modified chord diagrams to demonstrate the persistence of the active site waters for (C) TadA*0.1-RNA, (D) TadA*1.1-RNA, (E) TadA*0.1(L84F)-RNA, and (F) TadA*1.1(L84F)-RNA. (G) Reaction profile for the deprotonation of the activated water molecule the various TadA*-RNA systems.

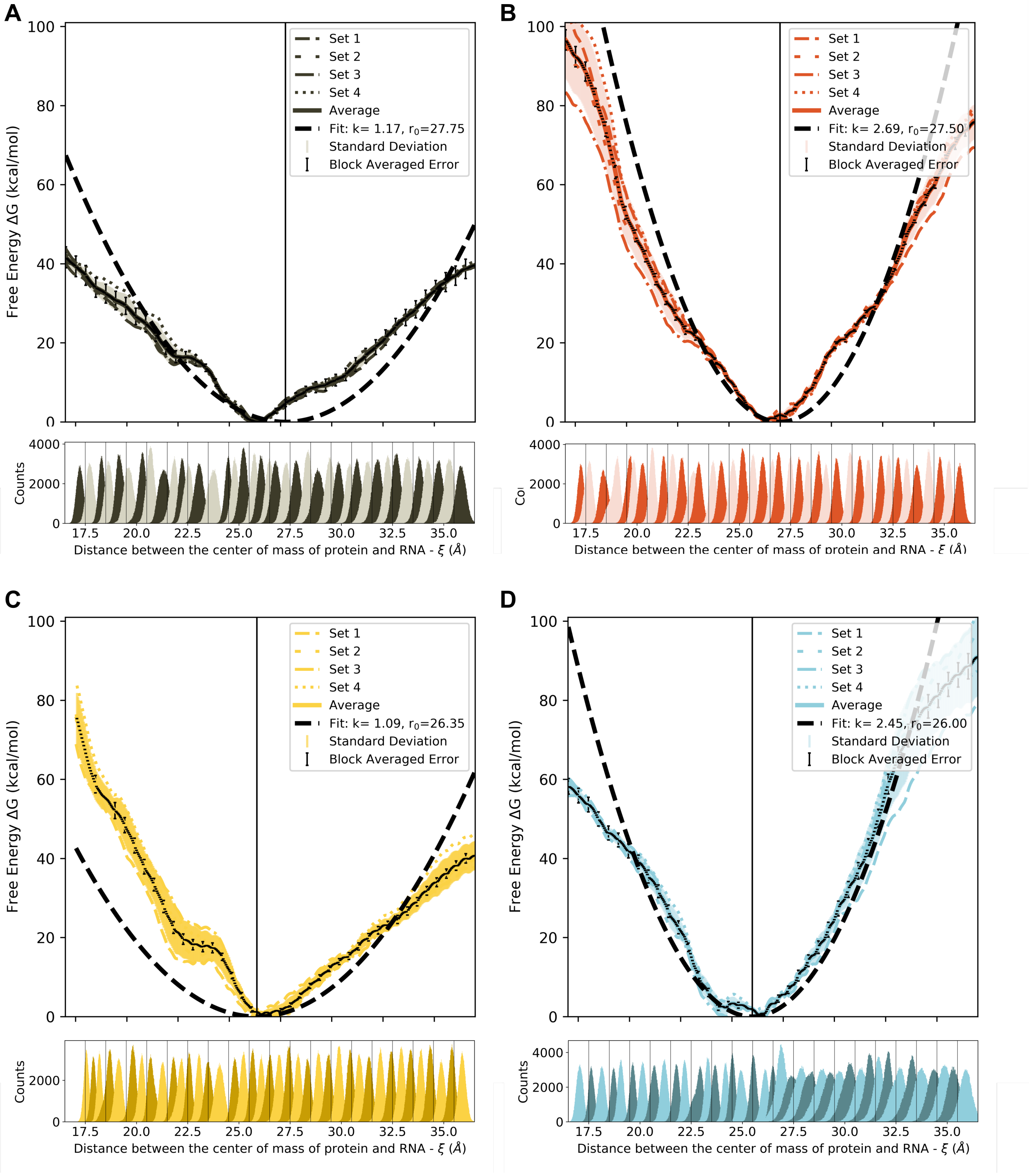

Figure S9: Umbrella sampling data and biased statistics. Individual pmfs associated to 4 independently conducted umbrella sampling simulations and the biased probability distributions (histograms) obtained from individual windows that stratify the $\xi$ space, for TadA*0.1-RNA (A), TadA*1.1-RNA (B), TadA*0.1(L84F)-RNA (C), and, TadA*1.1(L84F)-RNA (D) complex. In each pmf profile the shaded region indicated the standard deviation of the individual pmfs and the error bars are the error calculated using block averaging method for individual windows.

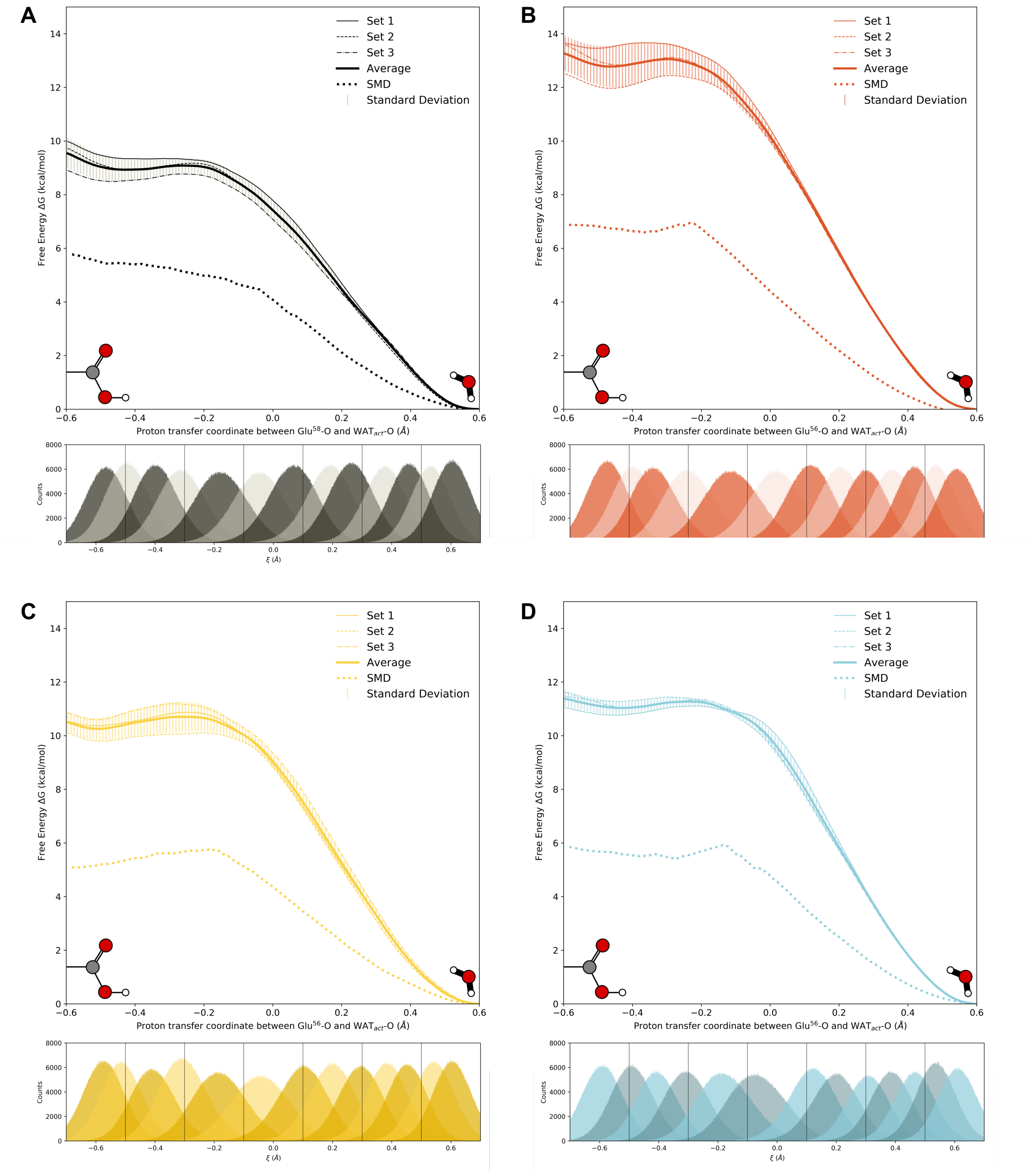

Figure S10: Steered MD,umbrella sampling data and biased statistics obtained from the QM/MM simulations of apo-TadA* systems. 3 individual pmfs that were used to compute the average pmf for the deprotonation reaction along the $\xi$ for (A) TadA*0.1, (B) TadA*1.1, (C) TadA*0.1(L84F), and (D) TadA*1.1(L84F). In each pmf profile the shaded region indicated the standard deviation of the average pmfs. The right extreme signifies that shared proton resides entirely on the activated water and the left extreme signifies that the shared proton resides entirely on the Glu$^{59}O$ atom.

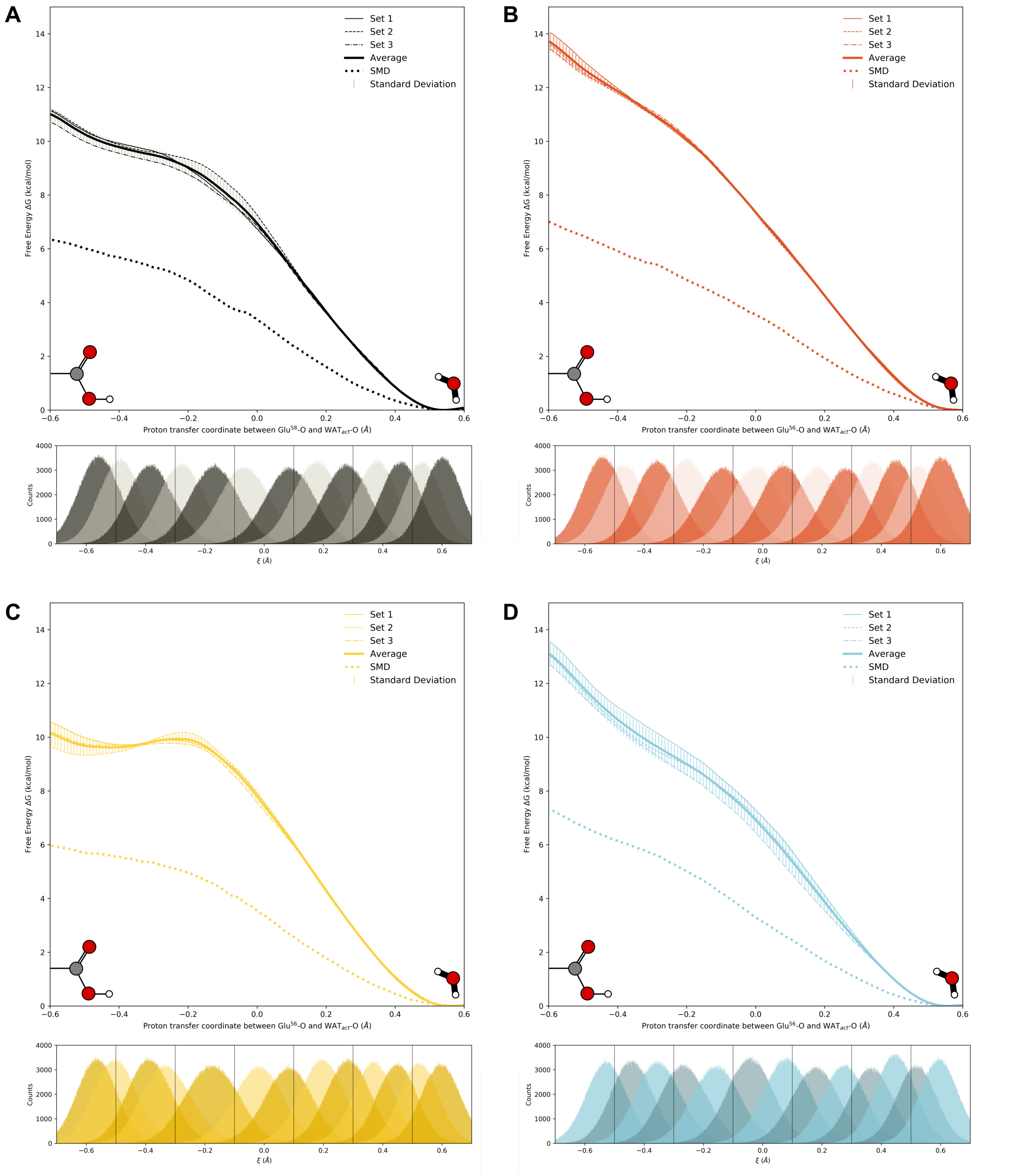

Figure S11: Steered MD, umbrella sampling data and biased statistics obtained from the QM/MM simulations of TadA*-RNA systems. 3 individual pmfs that were used to compute the average pmf for the deprotonation reaction along the $\xi$ for (A) TadA*0.1-RNA, (B) TadA*1.1-RNA, (C) TadA*0.1(L84F)-RNA, and (D) TadA*1.1(L84F)-RNA. In each pmf profile the shaded region indicated the standard deviation of the average pmfs. The right extreme signifies that shared proton resides entirely on the activated water and the left extreme signifies that the shared proton resides entirely on the $\mathrm{Glu}^{59}O$ atom.

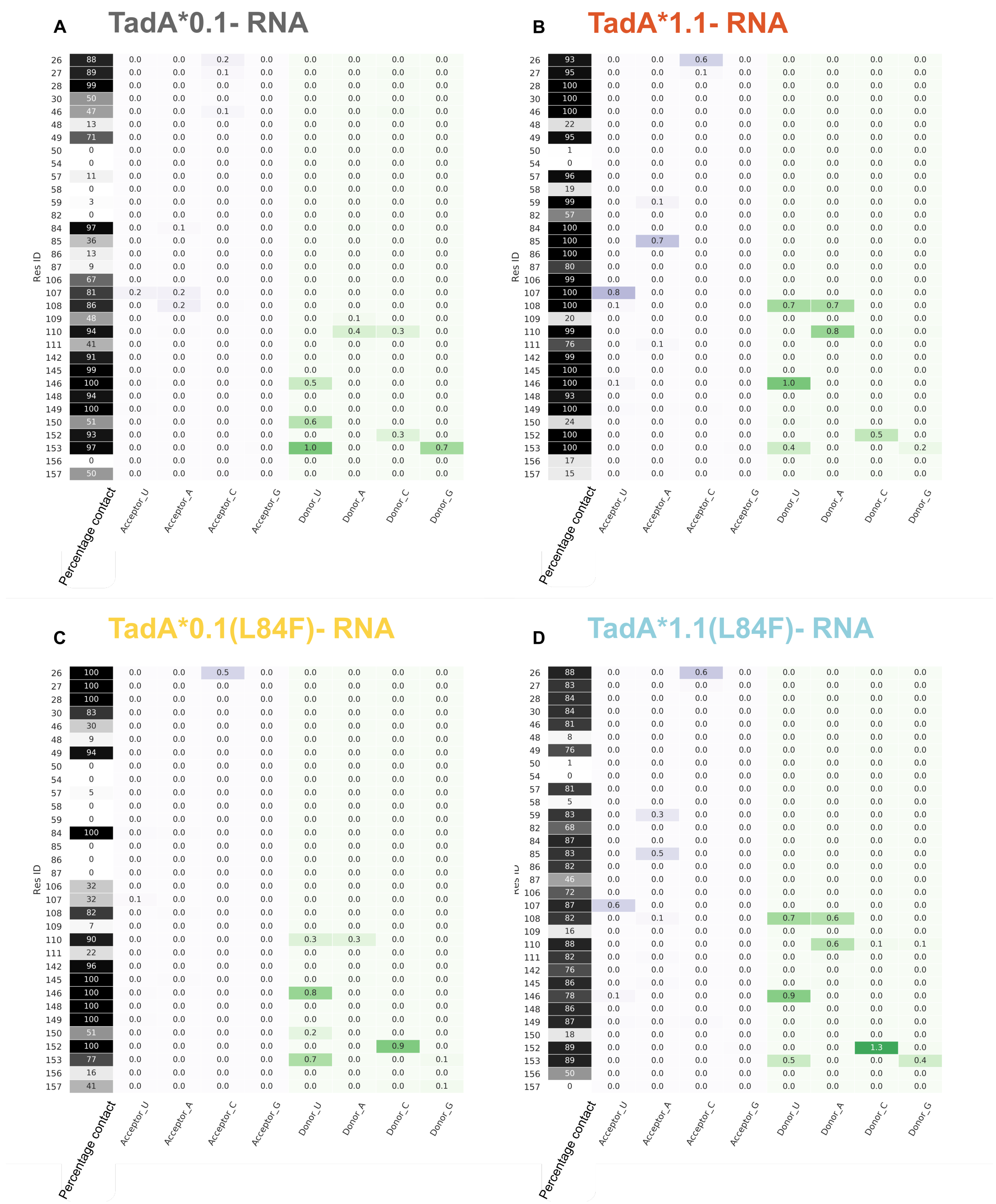

Figure S12: Percentage contact and the fractional H-bonding between the -UACG- consensus sequence nucleotides and the first interaction shell amino acids. For (A) TadA*0.1, (B) TadA*1.1, (C) TadA*0.1(L84F) and (D) TadA*1.1(L84F) in complex with RNA.The intensity of the colors in the columns signifies the magnitude of the percentage contact and the H-bonding strength.

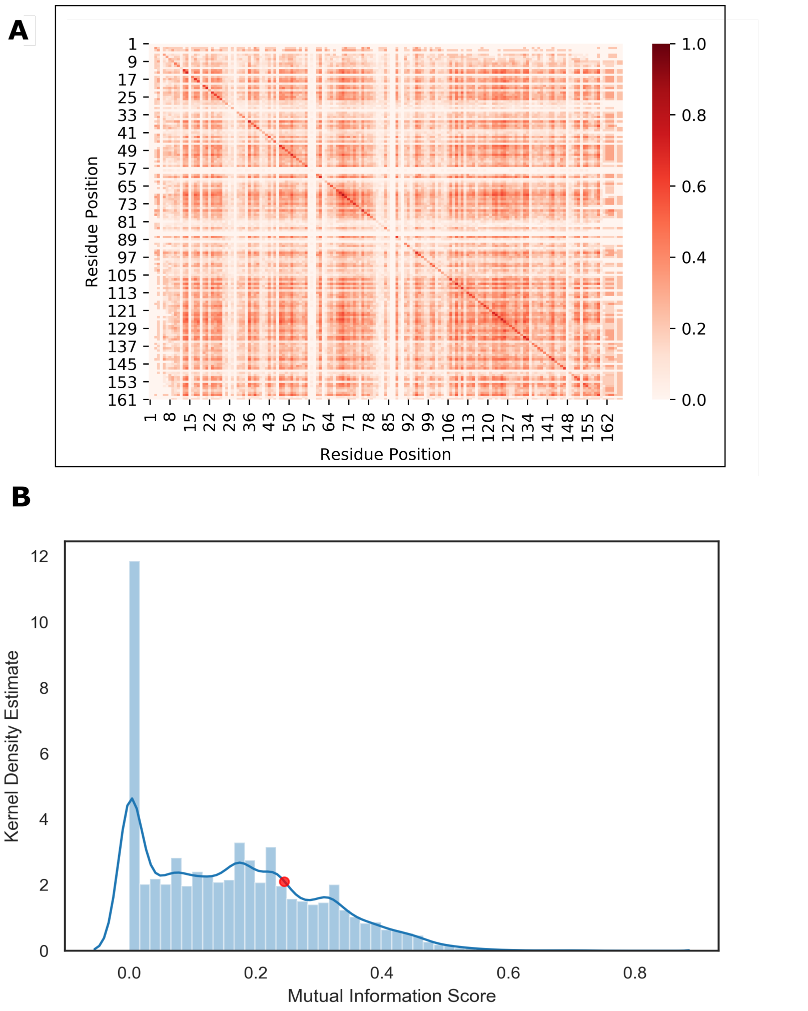

Figure S13: (A) Mutual information scores between pairs of residues for *E. coli* wtTadA. (B) Histogram showing the distribution of these mutual information scores. The value of the mutual information between residue 84 and 108 is indicated in red.

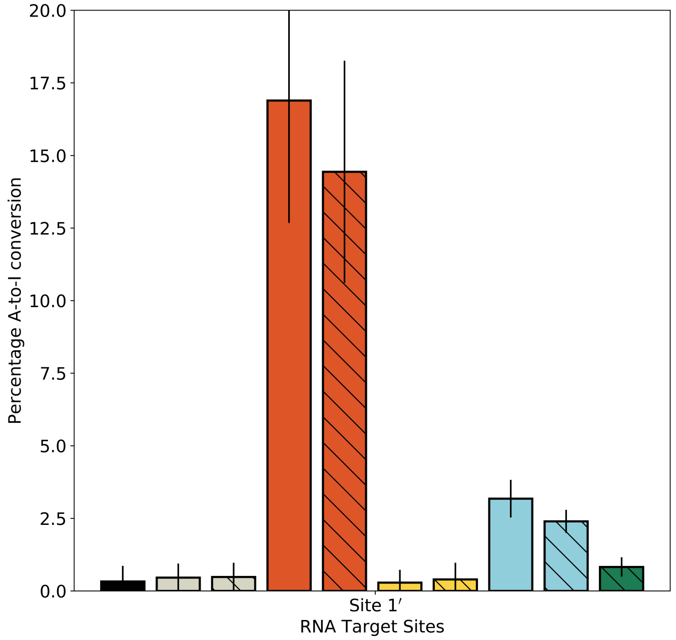

Figure S14: A-to-I base editing efficiencies in HEK293T cells by various ABE mutants at alternative RNA off-target site 1′. Values and error bars reflect the mean and SD of three independent biological replicates performed on different days.

#### 4 Supplementary Tables

| TadA*-RNA complex | TadA*0.1 | TadA*1.1 | TadA*0.1(L84F) | TadA*1.1(L84F) |
|---|---|---|---|---|
| $Zn^{2+}$-Target A distance (Å) | 4.25±0.64 | 3.34±0.32 | 6.65±0.61 | 3.60±0.31 |
| Res84 -Target A distance (Å) | 6.34±0.28 | 5.19±0.21 | 8.07±0.33 | 5.71±0.19 |

Table S1: Distances describing the conformational changes in the active site of Tad*–RNA complexes during the transition state formation. The $Zn^{+}2$-Target A distances are measured from the activated water to the C6 atom of target A base and the Res84-target A distances were measured by considering only the side change of residue 84 and the nucleobase of target A. The values here are the averages and S.D. corresponding to these distances in the umbrella sampling window with $\xi$=-0.5Å.

| Mutation | Frequency | Experimental Set |
|---|---|---|
| R13A | 3 | SECURE ABE |
| K20A | 0 | SECURE ABE |
| R21A | 0 | SECURE ABE |
| W23R | 1 | ABE 7.10, ABE 7.4 |
| W23L | 12 | ABE 7.8 |
| R*23A | 1 | SECURE ABE |
| E25A | 2 | SECURE ABE |
| R26A | 6 | SECURE ABE |
| E27A | 0 | SECURE ABE |
| V28G | 1 | SECURE ABE |
| V30G | 0 | SECURE ABE |
| H36L | 1 | ABE 5.3 |
| N46A | 0 | SECURE ABE |
| P48A | 0 | ABE 7.4 |
| P48S | 3 | ABE 6.4 |
| A**48G | 0 | SECURE ABE |
| I49A | 2 | SECURE ABE |
| R51L | 3 | ABE 5.3 |
| D53E | 0 | Zhuo et al. |
| A56G | 0 | SECURE ABE |
| E59A | 0 | Rees et al. |
| V82G | 0 | SECURE ABE |
| V82W | 0 | SECURE ABE |
| L84F | 0 | ABE 3.1 |
| E85A | 0 | SECURE ABE |
| P86A | 0 | SECURE ABE |
| C87A | 0 | SECURE ABE |
| C90A | 0 | SECURE ABE |
| A106V | 0 | ABE 1.2 |
| V***106G | 0 | SECURE ABE |
| V***106W | 0 | Rees et al. |
| D108N | 12 | ABE 1.1 |
| N^108A | 0 | SECURE ABE |
| A109G | 0 | SECURE ABE |
| T111A | 1 | SECURE ABE |
| H123Y | 0 | ABE 3.1 |
| A138G | 0 | SECURE ABE |
| A142L | 0 | ABE 4.3 |
| A142G | 2 | SECURE ABE |
| A143G | 1 | SECURE ABE |
| S146C | 0 | ABE 5.3 |
| D147Y | 0 | ABE 2.10 |
| F148A | 0 | SECURE ABE, Zhuo et al. |
| F149A | 0 | SECURE ABE |
| R152P | 0 | ABE 7.9 |
| P^^152A | 0 | SECURE ABE |
| E155V | 1 | ABE 2.10 |
| V^^^155G | 0 | SECURE ABE |
| V#155W | 0 | SECURE ABE |
| I156F | 0 | ABE 3.1 |
| A158N | 0 | ABE 5.3 |

Footnote
* From W23R in ABE 7.10
** From P48A in ABE 7.4
*** From A106V in ABE 1.2
^ From D108N in ABE 1.1
^^ From P152A in ABE 7.9
^^^ From E155V in ABE 2.10
### From E155V in ABE 2.10

Table S2: Analysis of the novelty of mutations in experiments by detecting frequency of prevalence across all hits in the filtered multiple sequence alignment output.

| System | Activated water | | Bridging water | |
|---|---|---|---|---|
| | Max. Persistence (ns) | Unique waters | Max. Persistence (ns) | Unique waters |
| TadA*0.1 | 347.7 | 3 | 525.9 | 2 |
| TadA*1.1 | 778.5 | 2 | 887.6 | 1 |
| TadA*0.1(L84F) | 375.4 | 10 | 162.0 | 8 |
| TadA*1.1(L84F) | 291.9 | 6 | 472.9 | 4 |
| TadA*0.1–RNA | 1000 | 1 | 1000 | 1 |
| TadA*1.1–RNA | 1000 | 1 | 1000 | 1 |
| TadA*0.1(L84F)–RNA | 642.2 | 3 | 667.6 | 5 |
| TadA*1.1(L84F)–RNA | 1000 | 1 | 1000 | 1 |

Table S3: Dynamics of the activated and bridging water molecules for various TadA* and TadA*–RNA systems.

#### 5 Supplementary Sequences

### ABE0.1-monomer

MKRTADGSEFESPKKKRKVSEVEFSHEYWMRHALTLAKRAWDEREVPVGAVLVHNNRVIGEGWNRPI
GRHDPTAHAEIMALRQGGLVMQNYRLIDATLYVTLEPCVMCAGAMIHSRIGRVVFGARDAKTGAAGS
LMDVLHHPGMNHRVEITEGILADECAALLSDFFRMRRQEIKAQKKAQSSTDSGGSSGGSSGSETPGT
SESATPESSGGSSGGSDKKYSIGLAIGTNSVGWAVITDEYKVPSKKFKVLGNTDRHSIKKNLIGALL
FDSGETAEATRLKRTARRRYTRRKNRICYLQEIFSNEMAKVDDSFFHRLEESFLVEEDKKHERHPIF
GNIVDEVAYHEKYPTIYHLRKKLVDSTDKADLRLIYLALAHMIKFRGHFLIEGDLNPDNSDVDKLFI
QLVQTYNQLFEENPINASGVDAKAILSARLSKSRRLENLIAQLPGEKKNGLFGNLIALSLGLTPNFK
SNFDLAEDAKLQLSKDTYDDDLDNLLAQIGDQYADLFLAAKNLSDAILLSDILRVNTEITKAPLSAS
MIKRYDEHHQDLTLLKALVRQQLPEKYKEIFFDQSKNGYAGYIDGGASQEEFYKFIKPILEKMDGTE
ELLVKLNREDLLRKQRTFDNGSIPHQIHLGELHAILRRQEDFYPFLKDNREKIEKILTFRIPYYVGP
LARGNSRFAWMTRKSEETITPWNFEEVVDKGASAQSFIERMTNFDKNLPNEKVLPKHSLLYEYFTVY
NELTKVKYVTEGMRKPAFLSGEQKKAIVDLLFKTNRKVTVKQLKEDYFKKIECFDSVEISGVEDRFN
ASLGTYHDLLKIIKDKDFLDNEENEDILEDIVLTLTLFEDREMIEERLKTYAHLFDDKVMKQLKRRR
YTGWGRLSRKLINGIRDKQSGKTILDFLKSDGFANRNFMQLIHDDSLTFKEDIQKAQVSGQGDSLHE
HIANLAGSPAIKKGILQTVKVVDELVKVMGRHKPENIVIEMARENQTTQKGQKNSRERMKRIEEGIK
ELGSQILKEHPVENTQLQNEKLYLYYLQNGRDMYVDQELDINRLSDYDVDHIVPQSFLKDDSIDNKV
LTRSDKNRGKSDNVPSEEVVKKMKNYWRQLLNAKLITQRKFDNLTKAERGGLSELDKAGFIKRQLVE
TRQITKHVAQILDSRMNTKYDENDKLIREVKVITLKSKLVSDFRKDFQFYKVREINNYHHAHDAYLN
AVVGTALIKKYPKLESEFVYGDYKVYDVRKMIAKSEQEIGKATAKYFFYSNIMNFFKTEITLANGEI
RKRPLIETNGETGEIVWDKGRDFATVRKVLSMPQVNIVKKTEVQTGGFSKESILPKRNSDKLIARKK
DWDPKKYGGFDSPTVAYSVLVVAKVEKGKSKKLKSVKELLGITIMERSSFEKNPIDFLEAKGYKEVK
KDLIIKLPKYSLFELENGRKRMLASAGELQKGNELALPSKYVNFLYLASHYEKLKGSPEDNEQKQLF
VEQHKHYLDEIIEQISEFSKRVILADANLDKVLSAYNKHRDKPIREQAENIIHLFTLTNLGAPAAFK
YFDTTIDRKRYTSTKEVLDATLIHQSITGLYETRIDLSQLGGDSGGSKRTADGSEFEPKKKRKV

### ABE0.1-dimer

MKRTADGSEFESPKKKRKVSEVEFSHEYWMRHALTLAKRAWDEREVPVGAVLVHNNRVIGEGWNRPI
GRHDPTAHAEIMALRQGGLVMQNYRLIDATLYVTLEPCVMCAGAMIHSRIGRVVFGARDAKTGAAGS
LMDVLHHPGMNHRVEITEGILADECAALLSDFFRMRRQEIKAQKKAQSSTDSGGSSGGSSGSETPGT
SESATPESSGGSSGGSSEVEFSHEYWMRHALTLAKRAWDEREVPVGAVLVHNNRVIGEGWNRPIGRH
DPTAHAEIMALRQGGLVMQNYRLIDATLYVTLEPCVMCAGAMIHSRIGRVVFGARDAKTGAAGSLMD
VLHHPGMNHRVEITEGILADECAALLSDFFRMRRQEIKAQKKAQSSTDSGGSSGGSSGSETPGTSES
ATPESSGGSSGGSDKKYSIGLAIGTNSVGWAVITDEYKVPSKKFKVLGNTDRHSIKKNLIGALLFDS
GETAEATRLKRTARRRYTRRKNRICYLQEIFSNEMAKVDDSFFHRLEESFLVEEDKKHERHPIFGNI
VDEVAYHEKYPTIYHLRKKLVDSTDKADLRLIYLALAHMIKFRGHFLIEGDLNPDNSDVDKLFIQLV
QTYNQLFEENPINASGVDAKAILSARLSKSRRLENLIAQLPGEKKNGLFGNLIALSLGLTPNFKSNF
DLAEDAKLQLSKDTYDDDLDNLLAQIGDQYADLFLAAKNLSDAILLSDILRVNTEITKAPLSASMIK
RYDEHHQDLTLLKALVRQQLPEKYKEIFFDQSKNGYAGYIDGGASQEEFYKFIKPILEKMDGTEELL
VKLNREDLLRKQRTFDNGSIPHQIHLGELHAILRRQEDFYPFLKDNREKIEKILTFRIPYYVGPLAR
GNSRFAWMTRKSEETITPWNFEEVVDKGASAQSFIERMTNFDKNLPNEKVLPKHSLLYEYFTVYNEL
TKVKYVTEGMRKPAFLSGEQKKAIVDLLFKTNRKVTVKQLKEDYFKKIECFDSVEISGVEDRFNASL
GTYHDLLKIIKDKDFLDNEENEDILEDIVLTLTLFEDREMIEERLKTYAHLFDDKVMKQLKRRRYTG
WGRLSRKLINGIRDKQSGKTILDFLKSDGFANRNFMQLIHDDSLTFKEDIQKAQVSGQGDSLHEHIA
NLAGSPAIKKGILQTVKVVDELVKVMGRHKPENIVIEMARENQTTQKGQKNSRERMKRIEEGIKELG
SQILKEHPVENTQLQNEKLYLYYLQNGRDMYVDQELDINRLSDYDVDHIVPQSFLKDDSIDNKVLTR
SDKNRGKSDNVPSEEVVKKMKNYWRQLLNAKLITQRKFDNLTKAERGGLSELDKAGFIKRQLVETRQ
ITKHVAQILDSRMNTKYDENDKLIREVKVITLKSKLVSDFRKDFQFYKVREINNYHHAHDAYLNAVV
GTALIKKYPKLESEFVYGDYKVYDVRKMIAKSEQEIGKATAKYFFYSNIMNFFKTEITLANGEIRKR
PLIETNGETGEIVWDKGRDFATVRKVLSMPQVNIVKKTEVQTGGFSKESILPKRNSDKLIARKKDWD
PKKYGGFDSPTVAYSVLVVAKVEKGKSKKLKSVKELLGITIMERSSFEKNPIDFLEAKGYKEVKKDL
IIKLPKYSLFELENGRKRMLASAGELQKGNELALPSKYVNFLYLASHYEKLKGSPEDNEQKQLFVEQ
HKHYLDEIIEQISEFSKRVILADANLDKVLSAYNKHRDKPIREQAENIIHLFTLTNLGAPAAFKYFD
TTIDRKRYTSTKEVLDATLIHQSITGLYETRIDLSQLGGDSGGSKRTADGSEFEPKKKRKV

### ABE1.1-monomer

MKRTADGSEFESPKKKRKVSEVEFSHEYWMRHALTLAKRAWDEREVPVGAVLVHNNRVIGEGWNRPI
GRHDPTAHAEIMALRQGGLVMQNYRLIDATLYVTLEPCVMCAGAMIHSRIGRVVFGAR**N**AKTGAAGS
LMDVLHHPGMNHRVEITEGILADECAALLSDFFRMRRQEIKAQKKAQSSTDSGGSSGGSSGSETPGT
SESATPESSGGSSGGSDKKYSIGLAIGTNSVGWAVITDEYKVPSKKFKVLGNTDRHSIKKNLIGALL
FDSGETAEATRLKRTARRRYTRRKNRICYLQEIFSNEMAKVDDSFFHRLEESFLVEEDKKHERHPIF
GNIVDEVAYHEKYPTIYHLRKKLVDSTDKADLRLIYLALAHMIKFRGHFLIEGDLNPDNSDVDKLFI
QLVQTYNQLFEENPINASGVDAKAILSARLSKSRRLENLIAQLPGEKKNGLFGNLIALSLGLTPNFK
SNFDLAEDAKLQLSKDTYDDDLDNLLAQIGDQYADLFLAAKNLSDAILLSDILRVNTEITKAPLSAS
MIKRYDEHHQDLTLLKALVRQQLPEKYKEIFFDQSKNGYAGYIDGGASQEEFYKFIKPILEKMDGTE
ELLVKLNREDLLRKQRTFDNGSIPHQIHLGELHAILRRQEDFYPFLKDNREKIEKILTFRIPYYVGP
LARGNSRFAWMTRKSEETITPWNFEEVVDKGASAQSFIERMTNFDKNLPNEKVLPKHSLLYEYFTVY
NELTKVKYVTEGMRKPAFLSGEQKKAIVDLLFKTNRKVTVKQLKEDYFKKIECFDSVEISGVEDRFN
ASLGTYHDLLKIIKDKDFLDNEENEDILEDIVLTLTLFEDREMIEERLKTYAHLFDDKVMKQLKRRR
YTGWGRLSRKLINGIRDKQSGKTILDFLKSDGFANRNFMQLIHDDSLTFKEDIQKAQVSGQGDSLHE
HIANLAGSPAIKKGILQTVKVVDELVKVMGRHKPENIVIEMARENQTTQKGQKNSRERMKRIEEGIK
ELGSQILKEHPVENTQLQNEKLYLYYLQNGRDMYVDQELDINRLSDYDVDHIVPQSFLKDDSIDNKV
LTRSDKNRGKSDNVPSEEVVKKMKNYWRQLLNAKLITQRKFDNLTKAERGGLSELDKAGFIKRQLVE
TRQITKHVAQILDSRMNTKYDENDKLIREVKVITLKSKLVSDFRKDFQFYKVREINNYHHAHDAYLN
AVVGTALIKKYPKLESEFVYGDYKVYDVRKMIAKSEQEIGKATAKYFFYSNIMNFFKTEITLANGEI
RKRPLIETNGETGEIVWDKGRDFATVRKVLSMPQVNIVKKTEVQTGGFSKESILPKRNSDKLIARKK
DWDPKKYGGFDSPTVAYSVLVVAKVEKGKSKKLKSVKELLGITIMERSSFEKNPIDFLEAKGYKEVK
KDLIIKLPKYSLFELENGRKRMLASAGELQKGNELALPSKYVNFLYLASHYEKLKGSPEDNEQKQLF
VEQHKHYLDEIIEQISEFSKRVILADANLDKVLSAYNKHRDKPIREQAENIIHLFTLTNLGAPAAFK
YFDTTIDRKRYTSTKEVLDATLIHQSITGLYETRIDLSQLGGDSGGSKRTADGSEFEPKKKRKV

### ABE1.1-dimer

MKRTADGSEFESPKKKRKVSEVEFSHEYWMRHALTLAKRAWDEREVPVGAVLVHNNRVIGEGWNRPI
GRHDPTAHAEIMALRQGGLVMQNYRLIDATLYVTLEPCVMCAGAMIHSRIGRVVFGARDAKTGAAGS
LMDVLHHPGMNHRVEITEGILADECAALLSDFFRMRRQEIKAQKKAQSSTDSGGSSGGSSGSETPGT
SESATPESSGGSSGGSSEVEFSHEYWMRHALTLAKRAWDEREVPVGAVLVHNNRVIGEGWNRPIGRH
DPTAHAEIMALRQGGLVMQNYRLIDATLYVTLEPCVMCAGAMIHSRIGRVVFGAR**N**AKTGAAGSLMD
VLHHPGMNHRVEITEGILADECAALLSDFFRMRRQEIKAQKKAQSSTDSGGSSGGSSGSETPGTSES
ATPESSGGSSGGSDKKYSIGLAIGTNSVGWAVITDEYKVPSKKFKVLGNTDRHSIKKNLIGALLFDS
GETAEATRLKRTARRRYTRRKNRICYLQEIFSNEMAKVDDSFFHRLEESFLVEEDKKHERHPIFGNI
VDEVAYHEKYPTIYHLRKKLVDSTDKADLRLIYLALAHMIKFRGHFLIEGDLNPDNSDVDKLFIQLV
QTYNQLFEENPINASGVDAKAILSARLSKSRRLENLIAQLPGEKKNGLFGNLIALSLGLTPNFKSNF
DLAEDAKLQLSKDTYDDDLDNLLAQIGDQYADLFLAAKNLSDAILLSDILRVNTEITKAPLSASMIK
RYDEHHQDLTLLKALVRQQLPEKYKEIFFDQSKNGYAGYIDGGASQEEFYKFIKPILEKMDGTEELL
VKLNREDLLRKQRTFDNGSIPHQIHLGELHAILRRQEDFYPFLKDNREKIEKILTFRIPYYVGPLAR
GNSRFAWMTRKSEETITPWNFEEVVDKGASAQSFIERMTNFDKNLPNEKVLPKHSLLYEYFTVYNEL
TKVKYVTEGMRKPAFLSGEQKKAIVDLLFKTNRKVTVKQLKEDYFKKIECFDSVEISGVEDRFNASL
GTYHDLLKIIKDKDFLDNEENEDILEDIVLTLTLFEDREMIEERLKTYAHLFDDKVMKQLKRRRYTG
WGRLSRKLINGIRDKQSGKTILDFLKSDGFANRNFMQLIHDDSLTFKEDIQKAQVSGQGDSLHEHIA
NLAGSPAIKKGILQTVKVVDELVKVMGRHKPENIVIEMARENQTTQKGQKNSRERMKRIEEGIKELG
SQILKEHPVENTQLQNEKLYLYYLQNGRDMYVDQELDINRLSDYDVDHIVPQSFLKDDSIDNKVLTR
SDKNRGKSDNVPSEEVVKKMKNYWRQLLNAKLITQRKFDNLTKAERGGLSELDKAGFIKRQLVETRQ
ITKHVAQILDSRMNTKYDENDKLIREVKVITLKSKLVSDFRKDFQFYKVREINNYHHAHDAYLNAVV
GTALIKKYPKLESEFVYGDYKVYDVRKMIAKSEQEIGKATAKYFFYSNIMNFFKTEITLANGEIRKR
PLIETNGETGEIVWDKGRDFATVRKVLSMPQVNIVKKTEVQTGGFSKESILPKRNSDKLIARKKDWD
PKKYGGFDSPTVAYSVLVVAKVEKGKSKKLKSVKELLGITIMERSSFEKNPIDFLEAKGYKEVKKDL
IIKLPKYSLFELENGRKRMLASAGELQKGNELALPSKYVNFLYLASHYEKLKGSPEDNEQKQLFVEQ
HKHYLDEIIEQISEFSKRVILADANLDKVLSAYNKHRDKPIREQAENIIHLFTLTNLGAPAAFKYFD
TTIDRKRYTSTKEVLDATLIHQSITGLYETRIDLSQLGGDSGGSKRTADGSEFEPKKKRKV

### ABE0.1(L84F)-monomer

MKRTADGSEFESPKKKRKVSEVEFSHEYWMRHALTLAKRAWDEREVPVGAVLVHNNRVIGEGWNRPI
GRHDPTAHAEIMALRQGGLVMQNYRLIDATLYVT**F**EPCVMCAGAMIHSRIGRVVFGARDAKTGAAGS
LMDVLHHPGMNHRVEITEGILADECAALLSDFFRMRRQEIKAQKKAQSSTDSGGSSGGSSGSETPGT
SESATPESSGGSSGGSDKKYSIGLAIGTNSVGWAVITDEYKVPSKKFKVLGNTDRHSIKKNLIGALL
FDSGETAEATRLKRTARRRYTRRKNRICYLQEIFSNEMAKVDDSFFHRLEESFLVEEDKKHERHPIF
GNIVDEVAYHEKYPTIYHLRKKLVDSTDKADLRLIYLALAHMIKFRGHFLIEGDLNPDNSDVDKLFI
QLVQTYNQLFEENPINASGVDAKAILSARLSKSRRLENLIAQLPGEKKNGLFGNLIALSLGLTPNFK
SNFDLAEDAKLQLSKDTYDDDLDNLLAQIGDQYADLFLAAKNLSDAILLSDILRVNTEITKAPLSAS
MIKRYDEHHQDLTLLKALVRQQLPEKYKEIFFDQSKNGYAGYIDGGASQEEFYKFIKPILEKMDGTE
ELLVKLNREDLLRKQRTFDNGSIPHQIHLGELHAILRRQEDFYPFLKDNREKIEKILTFRIPYYVGP
LARGNSRFAWMTRKSEETITPWNFEEVVDKGASAQSFIERMTNFDKNLPNEKVLPKHSLLYEYFTVY
NELTKVKYVTEGMRKPAFLSGEQKKAIVDLLFKTNRKVTVKQLKEDYFKKIECFDSVEISGVEDRFN
ASLGTYHDLLKIIKDKDFLDNEENEDILEDIVLTLTLFEDREMIEERLKTYAHLFDDKVMKQLKRRR
YTGWGRLSRKLINGIRDKQSGKTILDFLKSDGFANRNFMQLIHDDSLTFKEDIQKAQVSGQGDSLHE
HIANLAGSPAIKKGILQTVKVVDELVKVMGRHKPENIVIEMARENQTTQKGQKNSRERMKRIEEGIK
ELGSQILKEHPVENTQLQNEKLYLYYLQNGRDMYVDQELDINRLSDYDVDHIVPQSFLKDDSIDNKV
LTRSDKNRGKSDNVPSEEVVKKMKNYWRQLLNAKLITQRKFDNLTKAERGGLSELDKAGFIKRQLVE
TRQITKHVAQILDSRMNTKYDENDKLIREVKVITLKSKLVSDFRKDFQFYKVREINNYHHAHDAYLN
AVVGTALIKKYPKLESEFVYGDYKVYDVRKMIAKSEQEIGKATAKYFFYSNIMNFFKTEITLANGEI
RKRPLIETNGETGEIVWDKGRDFATVRKVLSMPQVNIVKKTEVQTGGFSKESILPKRNSDKLIARKK
DWDPKKYGGFDSPTVAYSVLVVAKVEKGKSKKLKSVKELLGITIMERSSFEKNPIDFLEAKGYKEVK
KDLIIKLPKYSLFELENGRKRMLASAGELQKGNELALPSKYVNFLYLASHYEKLKGSPEDNEQKQLF
VEQHKHYLDEIIEQISEFSKRVILADANLDKVLSAYNKHRDKPIREQAENIIHLFTLTNLGAPAAFK
YFDTTIDRKRYTSTKEVLDATLIHQSITGLYETRIDLSQLGGDSGGSKRTADGSEFEPKKKRKV

### ABE0.1(L84F)-dimer

MKRTADGSEFESPKKKRKVSEVEFSHEYWMRHALTLAKRAWDEREVPVGAVLVHNNRVIGEGWNRPI
GRHDPTAHAEIMALRQGGLVMQNYRLIDATLYVTLEPCVMCAGAMIHSRIGRVVFGARDAKTGAAGS
LMDVLHHPGMNHRVEITEGILADECAALLSDFFRMRRQEIKAQKKAQSSTDSGGSSGGSSGSETPGT
SESATPESSGGSSGGSSEVEFSHEYWMRHALTLAKRAWDEREVPVGAVLVHNNRVIGEGWNRPIGRH
DPTAHAEIMALRQGGLVMQNYRLIDATLYVT**F**EPCVMCAGAMIHSRIGRVVFGARDAKTGAAGSLMD
VLHHPGMNHRVEITEGILADECAALLSDFFRMRRQEIKAQKKAQSSTDSGGSSGGSSGSETPGTSES
ATPESSGGSSGGSDKKYSIGLAIGTNSVGWAVITDEYKVPSKKFKVLGNTDRHSIKKNLIGALLFDS
GETAEATRLKRTARRRYTRRKNRICYLQEIFSNEMAKVDDSFFHRLEESFLVEEDKKHERHPIFGNI
VDEVAYHEKYPTIYHLRKKLVDSTDKADLRLIYLALAHMIKFRGHFLIEGDLNPDNSDVDKLFIQLV
QTYNQLFEENPINASGVDAKAILSARLSKSRRLENLIAQLPGEKKNGLFGNLIALSLGLTPNFKSNF
DLAEDAKLQLSKDTYDDDLDNLLAQIGDQYADLFLAAKNLSDAILLSDILRVNTEITKAPLSASMIK
RYDEHHQDLTLLKALVRQQLPEKYKEIFFDQSKNGYAGYIDGGASQEEFYKFIKPILEKMDGTEELL
VKLNREDLLRKQRTFDNGSIPHQIHLGELHAILRRQEDFYPFLKDNREKIEKILTFRIPYYVGPLAR
GNSRFAWMTRKSEETITPWNFEEVVDKGASAQSFIERMTNFDKNLPNEKVLPKHSLLYEYFTVYNEL
TKVKYVTEGMRKPAFLSGEQKKAIVDLLFKTNRKVTVKQLKEDYFKKIECFDSVEISGVEDRFNASL
GTYHDLLKIIKDKDFLDNEENEDILEDIVLTLTLFEDREMIEERLKTYAHLFDDKVMKQLKRRRYTG
WGRLSRKLINGIRDKQSGKTILDFLKSDGFANRNFMQLIHDDSLTFKEDIQKAQVSGQGDSLHEHIA
NLAGSPAIKKGILQTVKVVDELVKVMGRHKPENIVIEMARENQTTQKGQKNSRERMKRIEEGIKELG
SQILKEHPVENTQLQNEKLYLYYLQNGRDMYVDQELDINRLSDYDVDHIVPQSFLKDDSIDNKVLTR
SDKNRGKSDNVPSEEVVKKMKNYWRQLLNAKLITQRKFDNLTKAERGGLSELDKAGFIKRQLVETRQ
ITKHVAQILDSRMNTKYDENDKLIREVKVITLKSKLVSDFRKDFQFYKVREINNYHHAHDAYLNAVV
GTALIKKYPKLESEFVYGDYKVYDVRKMIAKSEQEIGKATAKYFFYSNIMNFFKTEITLANGEIRKR
PLIETNGETGEIVWDKGRDFATVRKVLSMPQVNIVKKTEVQTGGFSKESILPKRNSDKLIARKKDWD
PKKYGGFDSPTVAYSVLVVAKVEKGKSKKLKSVKELLGITIMERSSFEKNPIDFLEAKGYKEVKKDL
IIKLPKYSLFELENGRKRMLASAGELQKGNELALPSKYVNFLYLASHYEKLKGSPEDNEQKQLFVEQ
HKHYLDEIIEQISEFSKRVILADANLDKVLSAYNKHRDKPIREQAENIIHLFTLTNLGAPAAFKYFD
TTIDRKRYTSTKEVLDATLIHQSITGLYETRIDLSQLGGDSGGSKRTADGSEFEPKKKRKV

Table S4: RNA sequences:

| RNA Site | Gene name | Amplicon |
|---|---|---|
| 1 | DNAJB1 | GCGCTACCACCCGGACAAGAACAAGGAGCCCGGCGCCG<br>AGGAGAAGTTCAAGGAGATCGCTGAGGCCT**A**CGACGTG<br>CTCAGCGACCCGCGCAAGCGCGAGATCTTCGACCGCTA<br>CGGGGAGGAAGGCCTAAAGGGGAGTGGCCCCAGTGGC<br>GGTAGCGGCGGTGGTGCCAATGGTACCTCTTTCAGCTA<br>CACATTCCATGGAGACCCTCATGCCATG |
| 2 | MTA2 | TCTGGCTTCAGGGATTCGTTCAAGCTCACAGCCAGCAGC<br>CAAGCGTCAGAAACTAAACCCAGCTGATGCCCCCAATCC<br>TGTGGTGTTTGTGGCCACAAAGGATACCAGGGCCCT**A**CG<br>GAAGGCTCTGACCCATCTGGAAATGCGGCGAGCTGCTCG<br>CCGACCCAACTTGCCCCTGAAGGTGAAGCCAACGCTGAT<br>TGCAGTGCGGCCCCCTGTCCCTCTACCTGCACCCTCACATC |
| 3 | PTBP2 | AGATTTTGGTAATTCCCCATTGCATCGTTTTAAGAAACCT<br>GGATCCAAAAATTTTCAAAACATTTTTCCTCCTTCTGCCAC<br>CCTTCACCTATCTAATATCCCTCCATCAGTAGCAGAAGAG<br>GATCT**A**CGAACACTGTTCGCTAACACTGGGGGCACTGTG<br>AAAGCATTTAAGTTTTTTCAAAGAGATCACAAAATGGCTCT<br>TCTTCAGATGGCAACAGTGGAAGAAGCTATTCAGG |
| 4 | SAP30BP | CAGAACCCCCTGGCAGATGTTCAAATCACTTGCAAGACA<br>AGATCCAGAAGCTTTATGAACGAAAGATAAAGGAGGGAA<br>TGGATATGAACTACATTATCCAAAGGAAGAAAGAATTTCG<br>GAACCCTAGCATCT**A**CGAGAAGCTGATCCAGTTCTGTGC<br>CATTGACGAGCTTGGCACCAACTACCCAAAGGATATGTTT<br>GATCCCCATGGCTGGTCTGAGGACTCCTACT |
| 5 | LCMT1 | ATTGCCAACACTCCTGATAGCTGAATGTGTGCTGGTTTAC<br>ATGACTCCAGAGCAGTCCGCAAACCTCCTGAAGTGGGCA<br>GCCAACAGTTTTGAGAGAGCCATGTTCATAAACT**A**CGAAC<br>AGGTGAACATGGGTGATCGGTTTGGGCAGATCATGATTG<br>AAAACCTGCGGAGACGCCAGTGTGACCTGGCGGGAGTG<br>GAGACCTGCAAGTC |
| 6 | SCAP | CCATTGACATTCGCCGGATGGAGCTAGCAGACCTGAACA<br>AGCGACTGCCCCCTGAGGCCTGCCTGCCCTCAGCCAAG<br>CCAGTGGGACAGCCAACGCGCT**A**CGAGCGGCAGCTGGC<br>TGTGAGGCCGTCCACACCCCACACCATCACGTTGCAGCC<br>GTCTTCCTTCCGAAACCTGCGGCTCCCCAAGAGGCTGCG<br>TGTTGTCTACTTC |
| gRNA sequence | | GGTATTACTGATATTGGTGGG |

Table S5: Primers:

| RNA Site | Primer 1 | Primer 2 |
|---|---|---|
| 1 | CATGGCATGAGGGTCTCCATGG | GCGCTACCACCCGGACAAG |
| 2 | GATGTGAGGGTGCAGGTAGAGGG | TCTGGCTTCAGGGATTCGTTCAAG |
| 3 | AGATTTTGGTAATTCCCCATTGCATCG | CCTGAATAGCTTCTTCCACTGTTGCC |
| 4 | CAGAACCCCCTGGCAGATGTTC | AGTAGGAGTCCTCAGACCAGCC |
| 5 | ATTGCCAACACTCCTGATAGCTGAATG | GACTTGCAGGTCTCCACTCCCG |
| 6 | GAAGTAGACAACACGCAGCCTCTTG | CCATTGACATTCGCCGGATGGAG |

I5 extension on primer 1s: ACACTCTTTCCCTACACGACGCTCTTCCGATCT

I7 extension on primer 2s: GACTGGAGTTCAGACGTGTGCTCTTCCGATCT
